# Transcriptional profiling of the murine airway response to acute ozone exposure

**DOI:** 10.1101/660316

**Authors:** Adelaide Tovar, Gregory J. Smith, Joseph M. Thomas, Jack R. Harkema, Samir N. P. Kelada

## Abstract

Exposure to ambient ozone (O_3_) pollution causes airway inflammation, epithelial injury, and decreased lung function. Long-term exposure is associated with increased mortality and exacerbations of respiratory conditions. While the adverse health effects of O_3_ exposure have been thoroughly described, less is known about the molecular processes that drive these outcomes. The aim of this study was to describe the cellular and molecular alterations observed in murine airways after exposure to either 1 or 2 ppm O_3_. After exposing adult, female C57BL/6J mice to filtered air, 1 or 2 ppm O_3_ for 3 hours, we assessed hallmark responses including airway inflammatory cell counts, epithelial permeability, cytokine secretion, and morphological alterations of the large airways. Further, we performed RNA-seq to profile gene expression in two critical tissues involved in O_3_ responses: conducting airways (CA) and airway macrophages (AM). We observed a concentration-dependent increase in airway inflammation and injury, and a large number of genes were differentially expressed in both target tissues at both concentrations of O_3_. Genes that were differentially expressed in CA were generally associated with barrier function, detoxification processes, and cellular proliferation. The differentially expressed genes in AM were associated with innate immune signaling, cytokine production, and extracellular matrix remodeling. Overall, our study has described transcriptional responses to acute O_3_ exposure, revealing both shared and unique gene expression patterns across multiple concentrations of O_3_ and in two important O_3_-responsive tissues. These profiles provide broad mechanistic insight into pulmonary O_3_ toxicity, and reveal a variety of targets for refined follow-up studies.

## Introduction

Ozone (O_3_) is a common urban air pollutant generated by photochemical reactions of primary pollutants including volatile organic compounds and nitrogen oxides. Exposure to ozone is associated with variety of adverse health outcomes, including increased mortality and cardiorespiratory morbidity (ITO 2005; DAY *et al*. 2017; MIROWSKY 2017). Upon inhalation, O_3_ causes pulmonary inflammation (ARIS 1993; DEVLIN 1996), decreased lung function (SCHELEGLE *et al*. 2009; KIM *et al*. 2011), and impaired epithelial barrier integrity (KEHRL *et al*. 1987; DEVLIN 1997). Together, these effects contribute to the incidence and exacerbation of chronic respiratory diseases including asthma and chronic obstructive pulmonary disease (COPD) (MCCONNELL 2002; KELLY AND FUSSELL 2011; GOODMAN *et al*. 2018; ZU *et al*. 2018). Regulatory measures have led to considerable improvements in air quality in recent decades; however, ground-level O_3_ concentrations are expected to rise with climate change-associated warming temperatures (BERNSTEIN AND RICE 2013; PFISTER 2014). Thus, ambient O_3_ exposure remains a critical public health concern.

Inhaled O_3_ readily reacts with cellular membranes and components of the lung lining fluid to generate bioactive mediators that induce oxidative stress, tissue injury, and innate immune signaling (PRYOR 1995; F.J. 2000). The lung epithelium and resident airway macrophages are the first two pulmonary cell types that encounter O_3_ and its reaction products, and previous work has established their critical roles in initiating and resolving O_3_-induced airway inflammation (SUNIL 2012; BAUER *et al*. 2015; MATHEWS 2015; SUNIL 2015). Further, O_3_ exposure causes damage to the airway epithelium and impairs macrophage phagocytic and efferocytic function, which can cause prolonged injury and inflammation (BECKER *et al*. 1991; GILMOUR *et al*. 1991; DEVLIN *et al*. 1994). Though previous studies have extensively described these processes (BROMBERG 2016), the exact molecular mechanisms that drive them have not been completely elucidated.

Using transcriptomic approaches is a powerful method to thoroughly probe responses to a given stimulus (SWEENEY *et al*. 2017). In the case of examining toxicant-induced responses, genomic profiling studies are useful for identifying markers of exposure and early effect and comprehensively describing a toxicant’s effects at the transcriptional level. Previous studies that investigated transcriptional responses in whole lung tissue and in inflammatory cells recruited to the lungs following O_3_ exposure have broadened our appreciation of O_3_-response pathways and mechanisms of toxicity, including the involvement of heat-shock proteins (*Hspa1*), extracellular matrix remodeling enzymes (*Mmp2*, *Mmp9*), and various proinflammatory signaling pathways (*Tnfr* family members, *Rela*), amongst others (GOHIL *et al*. 2003; NADADUR *et al*. 2005; KOOTER *et al*. 2007; BACKUS *et al*. 2010; BAUER *et al*. 2011; GABEHART *et al*. 2014; LEROY *et al*. 2015; VERHEIN 2015; WARD 2015; CIENCEWICKI *et al*. 2016).

Because whole lung tissue is composed of a complex mixture of many (perhaps > 40) cell types (FRANKS 2008), bulk transcriptomics may reflect only the most marked alterations in gene expression; more subtle effects, including those that are tissue-specific, may be obscured. Therefore, approaches that focus on specific target tissues and/or cell types are required to resolve heterogeneity in gene expression responses across individual compartments and facilitate their clearer interpretation. To this end, we designed a study to examine transcriptional responses in the airway epithelium and airway macrophages after exposure to multiple concentrations of O_3_.

We exposed adult, female C57BL/6J mice to filtered air, 1 or 2 ppm O_3_ for three hours. Twenty-one hours later, we evaluated hallmark pulmonary inflammation and injury responses to O_3_ exposure in conjunction with gene expression profiling of the conducting airways (CA) and airway macrophages (AM). We found that the expression of a large number of genes was altered in a concentration- and tissue-dependent manner, including many genes not reported previously as O_3_-responsive.

## Materials and Methods

### Animals

Adult (8 weeks of age) female C57BL6/J mice were purchased from the Jackson Laboratory (Bar Harbor, Maine). All animals were housed in groups of three or more in polycarbonate cages on ALPHA-Dri bedding (Shepard), under normal 12-hour light/dark cycles with *ad libitum* food (Envigo 2929) and water. All experiments were approved by the Institutional Animal Care and Use Committee at the University of North Carolina at Chapel Hill.

### Ozone exposure

Mice were exposed to filtered air, 1 or 2 ppm ozone (O_3_) for three hours in individual wire-mesh chambers without access to food or water, as described previously (SMITH *et al*. 2019). Exposures at each concentration were performed on separate days to ensure that the exposure time (9 am-12 pm) was kept consistent for each exposure group, and the filtered air exposure group was split in half and one half was used as the matched controls for the 1 and 2 ppm exposure groups. At the end of the three hour exposure period, mice were returned to their normal housing.

### Phenotyping

#### Lung phenotyping

Eleven mice per treatment group were used for lung phenotyping and gene expression analyses. Twenty-one hours after exposure, mice were anesthetized (2 g/kg urethane) and sacrificed by exsanguination via the abdominal aorta/inferior vena cava. Bronchoalveolar lavage (BAL) was performed by cannulating the trachea and instilling phosphate-buffered saline supplemented with cOmplete protease inhibitor cocktail (Roche) (0.5 ml x 1, 1 ml x 1). The right upper and middle lobes were snap-frozen in liquid nitrogen and stored at -80°C. The remaining lobes and a section of the trachea were transferred to RNA*later* solution (Sigma Aldrich) and stored at 4°C until microdissection and RNA extraction. The recovered BAL fluid was centrifuged at 2,000xg for 10 minutes. The supernatant from the first fraction was saved and stored at -80*°*C as two aliquots for protein and cytokine analysis. Pellets from both fractions were pooled, washed once in red blood cell lysis buffer, and centrifuged again as previously. Pellets were resuspended in 500 uL of HBSS, a 20 uL cell suspension was added to 20 uL of Trypan blue solution, and total numbers of viable cells were determined by counting on a hemacytometer. A 100 uL aliquot of the cell suspension was used to prepare cytospin slides. The remaining cell suspension was plated in FBS-containing RPMI-1640 and rested in a cell culture incubator for 4 hours to enrich for airway macrophages (adherent cells).

#### Protein and cytokine analysis

BAL aliquots for total protein analysis were thawed on ice. Total protein was measured using the Qubit total protein quantification kit and Qubit 2.0 fluorometer (Thermo Scientific). Cytokines were measured using a Milliplex immunoassay kit (Millipore) on a Bio-Rad Bio-Plex 200 multiplex suspension array system.

#### Histology

Mice exposed exclusively for histological analyses (n = 5 per treatment group) were not subjected to BAL to avoid eliminating infiltrating immune cells and disrupting the natural architecture of the lungs. Mice were sacrificed as described in the previous section. Lung tissue was inflated *in situ* with 10% neutral-buffered formalin (NBF) through the tracheal cannula at 25 cm of static fluid pressure. The lungs were removed and immersed in 10% NBF for 24 hours overnight, followed by washing and dehydration in 70% ethanol. After fixation, lungs were embedded in paraffin and sectioned. Paraffin sections were stained with hematoxylin and eosin, and were immunostained with antibodies specific for FOXJ1 (Abcam, cat# ab235445, clone EPR21874, 1:1000 dilution) or CCSP (Abcam, cat#ab40873, 1:2000 dilution).

### RNA isolation and sequencing

#### Airway macrophage (AM) gene expression analysis

AM enriched from the BAL cell suspension were lysed directly with 350 uL of RLT buffer (Qiagen). Lysed AM were combined to create four pools per treatment group, and total RNA was isolated using the Qiagen RNeasy Micro Kit. Libraries were prepared using the Takara Bio Low Input SMARTer Stranded RNA-Seq Library Kit. Libraries were pooled and sequenced on one lane of an Illumina HiSeq 4000 to generate single-end, 50-bp reads.

#### Conducting airway (CA) gene expression analysis

Four lung samples were selected from each treatment group, representative of the four pools of AMs. CAs were isolated from lungs preserved in RNA*later* using a previously published method (BAKER *et al*. 2004). We isolated total RNA using the Qiagen RNeasy Mini Kit. Libraries were prepared from poly A-enriched RNA using the Kapa Stranded RNA-Seq Library Kit. Libraries were pooled and sequenced on one lane of an Illumina HiSeq 4000 to generate single-end, 50-bp reads.

### Gene expression analysis

Given the differences in the library preparation methods, analysis of CA and AM samples were performed separately rather than jointly, and only post hoc comparisons between the two compartments were made.

#### Sequence alignment and transcript quantification

After sequencing, reads were de-multiplexed and deposited as fastq files. Reads were aligned to the C57BL/6J [mm9, GENCODE release M18] reference genome using STAR v.2.6.0a (DOBIN *et al*. 2013). Transcripts were quantified with Salmon v0.9.1 using default parameters (PATRO 2017).

#### Differential expression analysis

tximport was used to import and summarize Salmon quantification files. Differentially expressed genes were identified using the standard differential expression analysis in DESeq2 (LOVE *et al*. 2014), extracting pairwise comparisons between each treatment group within a tissue compartment. A gene was considered differentially expressed if the absolute log_2_ fold change was greater than 1 and the Benjamini-Hochberg adjusted p-value (false discovery rate (FDR)) was less than 0.05.

#### Gene expression categorization

We binned genes into specific trend categories using the following conditions:

| Trend | Formula |
| --- | --- |
| **Positive monotonic** | $LFC_{1,FA} > 1 \ \& \ p\text{-val} < 0.05, LFC_{2,FA} > 1 \ \& \ p\text{-val} < 0.05, LFC_{2,1} > 1 \ \& \ p\text{-val} < 0.05$ |
| **Positive threshold A** | $LFC_{1,FA} > 1 \ \& \ p\text{-val} < 0.05, LFC_{2,FA} > 1 \ \& \ p\text{-val} < 0.05, \text{abs}(LFC_{2,1}) < 0.3 \ \& \ p\text{-val} > 0.1$ |
| **Positive threshold B** | $\text{abs}(LFC_{1,FA}) < 0.3 \ \& \ p\text{-val} > 0.1, LFC_{2,FA} > 1 \ \& \ p\text{-val} < 0.05, LFC_{2,1} > 1 \ \& \ p\text{-val} < 0.05$ |
| **Negative monotonic** | $LFC_{1,FA} < -1 \ \& \ p\text{-val} < 0.05, LFC_{2,FA} < -1 \ \& \ p\text{-val} < 0.05, LFC_{2,1} < -1 \ \& \ p\text{-val} < 0.05$ |
| **Negative threshold A** | $LFC_{1,FA} < -1 \ \& \ p\text{-val} < 0.05, LFC_{2,FA} < -1 \ \& \ p\text{-val} < 0.05, \text{abs}(LFC_{2,1}) < 0.3 \ \& \ p\text{-val} > 0.1$ |
| **Negative threshold B** | $\text{abs}(LFC_{1,FA}) < 0.3 \ \& \ p\text{-val} > 0.1, LFC_{2,FA} < -1 \ \& \ p\text{-val} < 0.05, LFC_{2,1} < -1 \ \& \ p\text{-val} < 0.05$ |
| **Peak** | $LFC_{1,FA} > 1 \ \& \ p\text{-val} < 0.05, \text{abs}(LFC_{2,FA}) < 0.3 \ \& \ p\text{-val} > 0.1, LFC_{2,1} < -1 \ \& \ p\text{-val} < 0.05$ |
| **Trough** | $LFC_{1,FA} < -1 \ \& \ p\text{-val} < 0.05, \text{abs}(LFC_{2,FA}) < 0.3 \ \& \ p\text{-val} > 0.1, LFC_{2,1} > 1 \ \& \ p\text{-val} < 0.05$ |

LFC: log<sub>2</sub> fold-change, 1.FA: 1 ppm versus filtered air, 2.FA: 2 ppm versus filtered air, 2.1: 1 ppm versus filtered air, abs: absolute value

#### Pathway analysis

We used Gene Set Variation Analysis (GSVA) (HÄNZELMANN 2013) to identify pathways that were differentially expressed in CA and AM due to O_3_ exposure. The variance-stabilized transformation of the gene expression count matrix from DESeq2 was used as input, and gene set libraries were downloaded from the Enrichr website (date: 01/22/2019, URLs in Supplemental Table 10) (KULESHOV *et al*. 2016). Resulting pathway enrichment scores were tested for differential enrichment using limma (RITCHIE *et al*. 2015). Gene sets were considered significantly enriched if the Benjamini-Hochberg adjusted p-value was less than 0.05.

#### Literature search and meta-analysis

PubMed and Google Scholar were searched for studies that evaluated gene expression responses to acute O_3_ exposure. We excluded studies that used lower (i.e., < 0.5 ppm) or higher (> 2 ppm) doses of O_3_ than our study, or chronic exposure. Some studies included in this analysis included genetically engineered mice. For these studies, only data collected from wild-type animals were used. For studies that published expression of microarray probes that had not been assigned final gene annotations (i.e., were published with accession numbers only), the genes were systematically assigned symbol names using the DAVID Gene ID Conversion Tool or manually assigned symbol names using Ensembl. From our study, we only used genes that were differentially expressed in the 1 ppm versus filtered air. A hypergeometric test (using a total background gene set size of 20,000) was performed to determine the significance of overlap between our results and a given study’s results, and we used all reported significant DEGs from the study, regardless of the magnitude of fold-change or the directionality of effect. The input lists of genes and all comparisons with p-values are included in the Supplemental Material. We also evaluated how consistently genes were considered differentially expressed across all 3 published studies and ours using the “vote counting” method of meta-analysis (RAMASAMY *et al*. 2008) in which we simply tallied the number of studies in which a gene was considered differentially expressed.

### Statistical analysis

For analysis presented in Figures 1 and 2, all raw data were subjected to Box-Cox power transformations to reduce heteroscedasticity and to conform to a normal distribution. Subsequently, we performed ANOVA and pairwise t-tests in R (version 3.5.3), and tests were considered significant if the resulting p-value was < 0.05.

**Figure 1.**
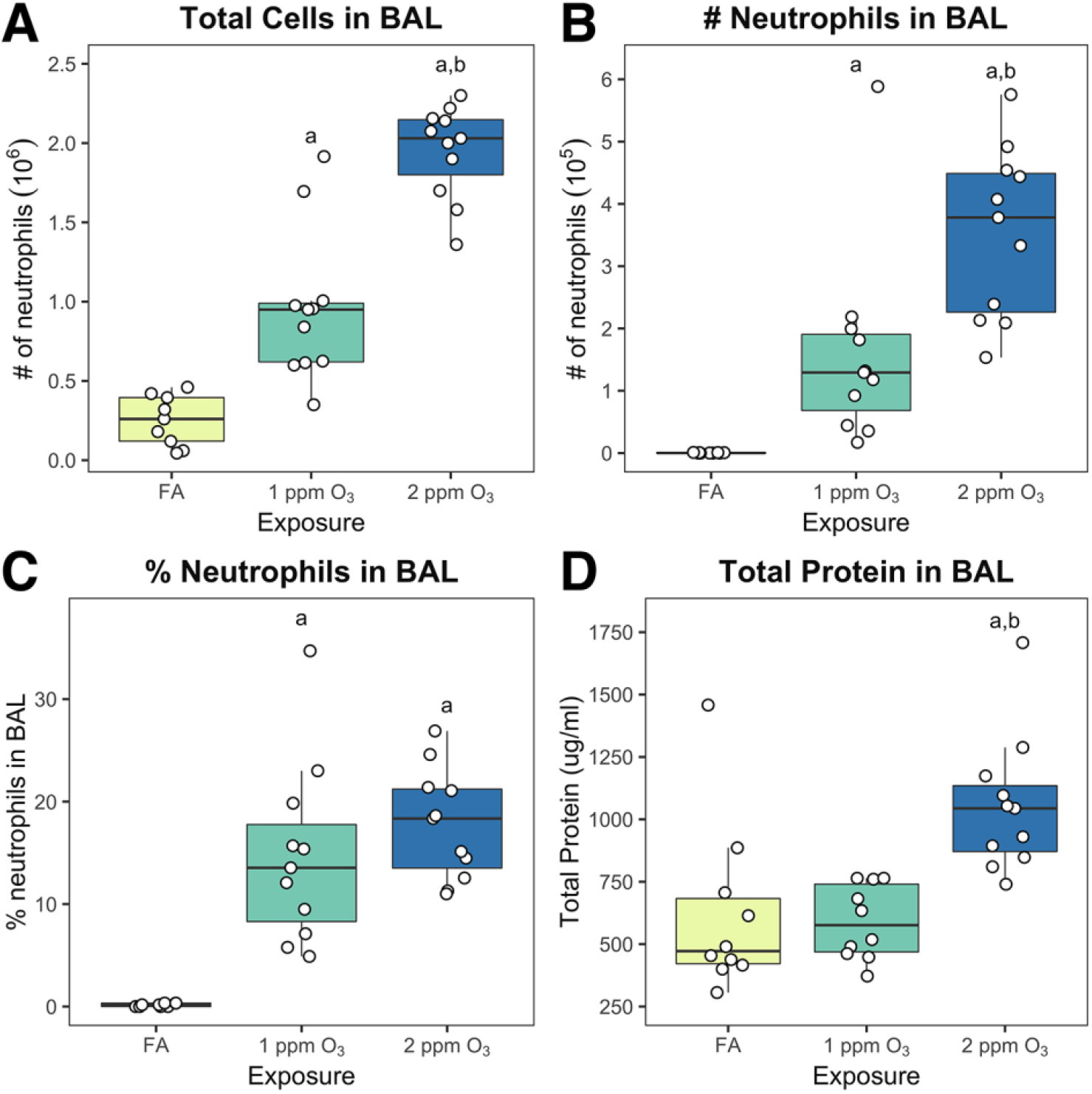
O_3_ exposure induces inflammation and injury in C57BL/6J mice. Eight week-old female C57BL/6J mice were exposed to filtered air (FA), 1 or 2 ppm O_3_ for 3 hours, sacrificed 21 hours subsequently, and cell types, cell numbers, and total protein concentration were measured in bronchoalveolar lavage fluid. (A) Total cell number (10^6^), (B) neutrophil number (10^5^), and (C) % neutrophils were measured by differential cell counting. (D) Total protein in BAL was measured using a fluorometric Qubit assay. Results are displayed as box-and-whisker plots depicting the distribution of the data as the minimum, first quartile, median, third quartile, and maximum with all points overlaid. (n=11/treatment group; a: p<0.05 compared to FA group, b: p<0.05 compared to 1 ppm group)

**Figure 2.**
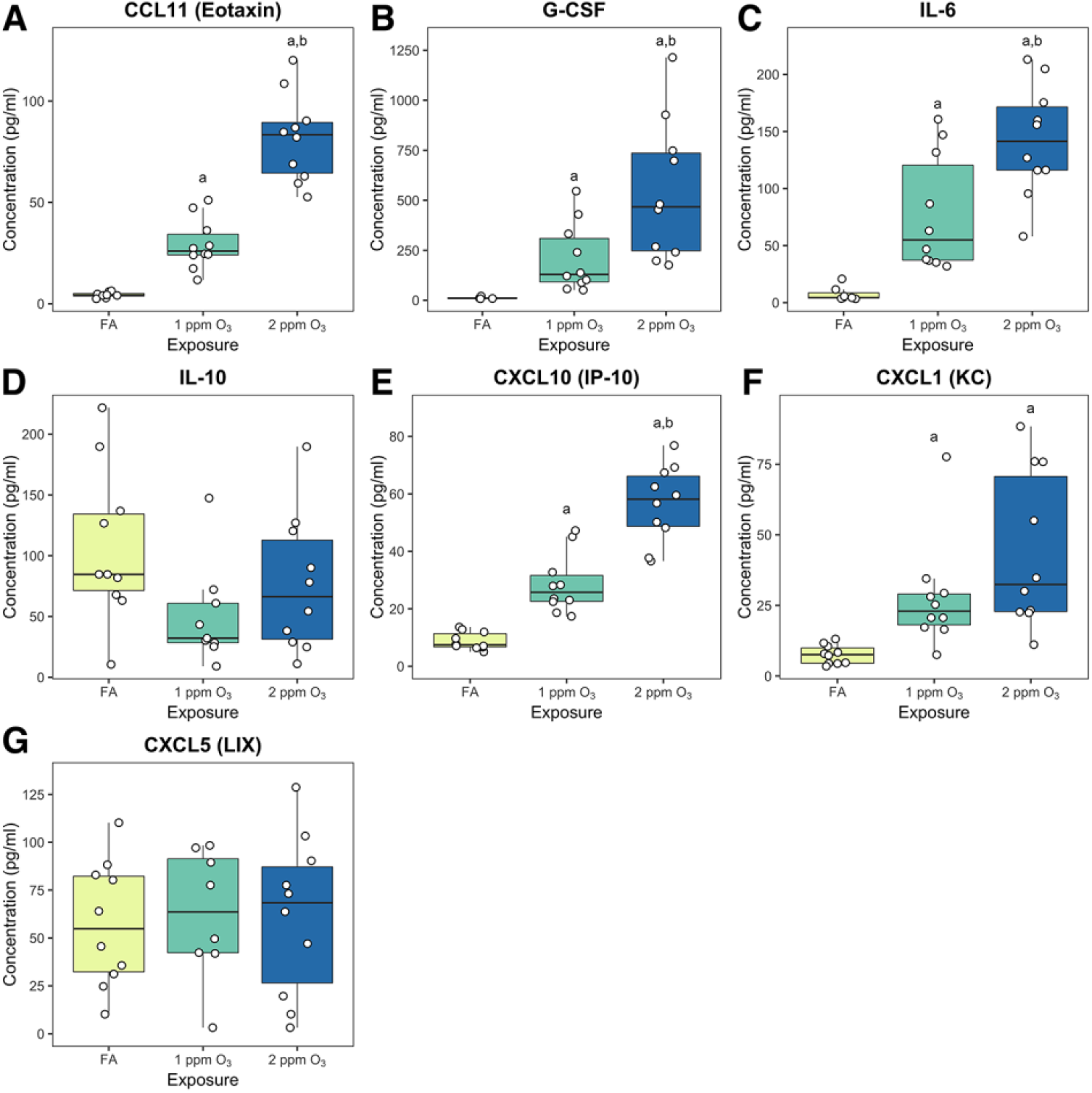
Respiratory cytokine and chemokine secretion are altered following O_3_ exposure. A multiplex cytokine detection assay was used to measure the secretion of (A) CCL11 (eotaxin), (B) G-CSF, (C) IL-6, (D), IL-10, (E) CXCL10 (IP-10), (F) CXCL1 (KC), and (G) CXCL5 (LIX). Results are displayed as box-and-whisker plots depicting the distribution of the data as the minimum, first quartile, median, third quartile, and maximum with all points overlaid. (n=10/treatment group were assayed, and points below limited of detection were excluded from analysis; a: p<0.05 compared to FA group, b: p<0.05 compared to 1 ppm group)

## Results

### Ozone (O_3_) induces inflammation, injury and altered cytokine production at two concentrations

We exposed 8 week-old, female C57BL/6J mice to filtered air (FA), 1 or 2 ppm O_3_ for 3 hours. Twenty-one hours after exposure, the total number of cells and neutrophils (as well as the percentage of neutrophils) in bronchoalveolar lavage (BAL) were significantly increased in mice exposed to either 1 or 2 ppm O_3_ compared to mice exposed to FA (Fig. 1A-C). Additionally, total cells and total number of neutrophils in BAL were significantly increased in mice exposed to 2 ppm O_3_ compared to those exposed to 1 ppm O_3_ (Fig. 1A-B), providing evidence of a roughly linear concentration-response relationship for these endpoints. In contrast to inflammation, lung injury (as measured by total protein in BAL fluid) was only apparent after exposure to 2 ppm O_3_, indicating a threshold type effect.

To further characterize airway inflammatory responses, we measured secretion of a panel of chemo- and cytokines. We observed concentration-dependent increases in canonical O_3_-responsive cytokines including eotaxin (CCL11), G-CSF, IL-6, IP-10 (CXCL10), and KC (CXCL1) (Fig. 2A-C, E-F) (LU *et al*. 2006). Contrary to expectation, we did not detect an increase in IL-10 or LIX (CXCL5), two cytokines that have been previously implicated in O_3_ responses (LU *et al*. 2006). IL-1β, IL-10, IL-12p70, MCP-1 (CCL2), MIP-1α (CCL3), MIP-1β (CCL4), MIP-2 (CXCL2), and TNF-α were also included in our multiplex panel, but were not detected above the minimum threshold (data not shown).

### Ozone induces epithelial injury and morphological changes to the airways

We performed histopathological analysis to assess airway morphological changes induced by O_3_ exposure (Fig. 3). At both concentrations of O_3_, we observed denudation of the epithelium and exfoliation of ciliated cells, both of which were more severe in mice exposed to 2 ppm O_3_ (Fig. 3A). Using immunohistochemical approaches, we found a marked loss of FOXJ1 protein, a marker of airway ciliated cells, in the axial airways of mice exposed to 1 and 2 ppm O_3_ (Fig. 3B). In contrast, while there was an appreciable change in the pattern of the secretory cell marker CCSP, its expression remained high in airway epithelial cells, which may be attributable to new, proliferating cells (Fig. 3C).

**Figure 3.**
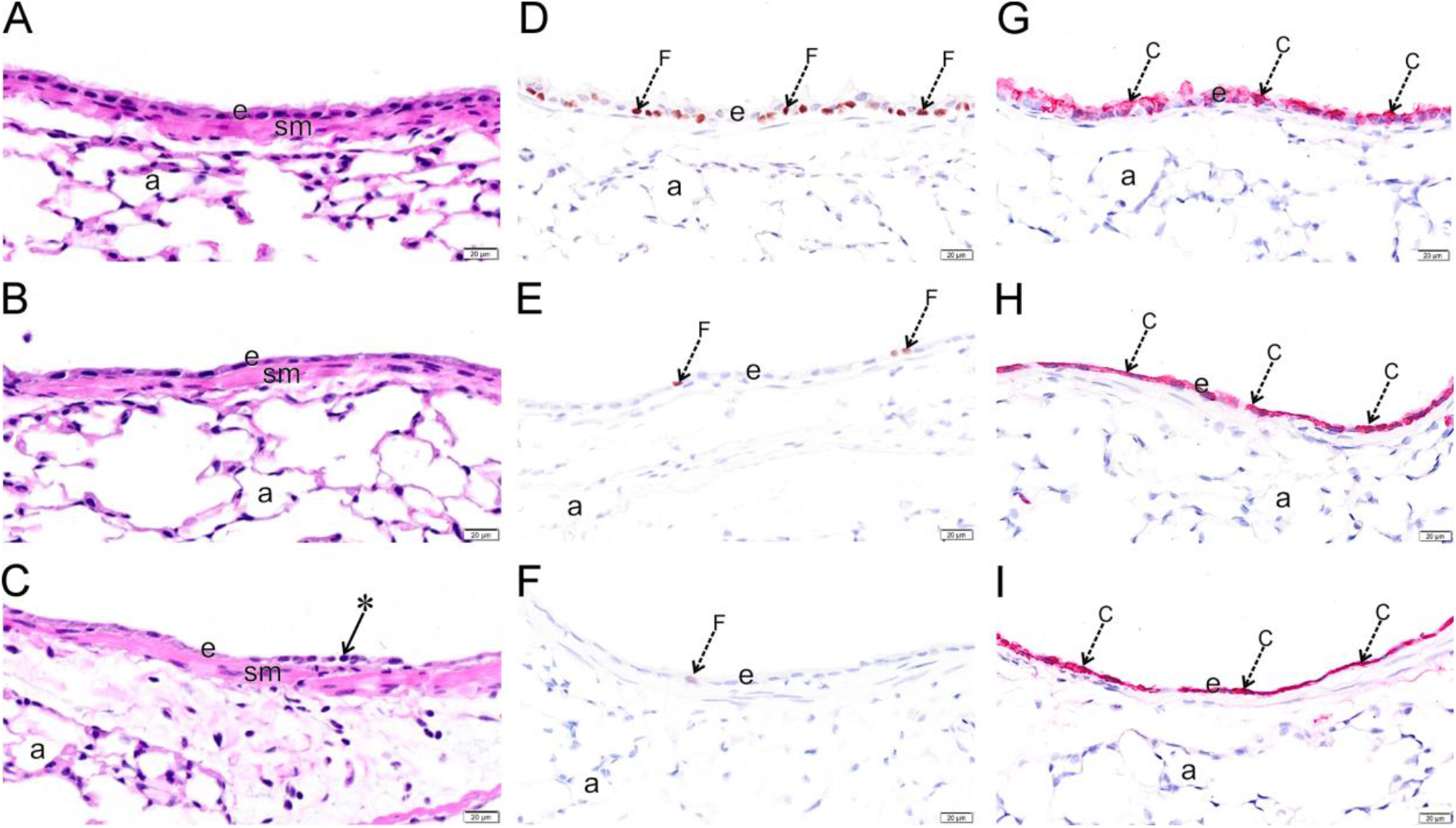
O_3_ exposure causes epithelial damage in the upper airways. Light photomicrographs of the respiratory epithelium lining the axial airway of the left lung lobe from mice exposed to 0 (A, D, G), 1 (B, E, H) or 2 ppm (C, F, I) ozone. Tissues were histochemically stained with hematoxylin and eosin (A, B, C) or immunohistochemically stained for (D, E, F) FOXJ1 or (G, H, I) club cell secretory protein (CCSP). e, respiratory epithelium; sm, airway smooth muscle; a, alveolar parenchyma; arrow *, exfoliating epithelium, arrows F (brown chromogen), FOXJ1; arrow C (red chromogen), Club Cell Secretory Protein.

### Marked transcriptional alterations in both conducting airways (CA) and airway macrophages (AM) following O_3_ exposure

We sought to examine transcriptional activity in CA tissue and AM twenty-one hours after FA, 1 ppm or 2 ppm O_3_ exposure. Here, we define CA as the large airway tree beginning at the trachea and ending at the terminal bronchioles, which captures 4 generations of branching airways (BAKER *et al*. 2004) and is highly susceptible to O_3_ toxicity (PLOPPER 1994). We use the term airway macrophages to describe the enriched population of adherent cells we collected from BAL. In O_3_-exposed mice, this likely includes both resident macrophages (TIGHE 2011) (including alveolar macrophages (SUNIL 2012; BIRUKOVA *et al*. 2019)) and recruited monocyte-derived macrophages (FRANCIS *et al*. 2017a).

RNA-seq was performed on 12 CA samples and 12 pooled AM samples (n=4/treatment group), and yielded an average of 27 million reads per sample, after preprocessing. Of these, an average of 81% of reads in CA and 75% of reads in AM were uniquely mapped, without any evidence of differential alignment across treatment groups. Additional mapping statistics are provided in Supplemental Table 1. Principal components analysis (PCA) of normalized gene expression profiles revealed a clear separation between the three treatment groups in both CA and AM samples (Fig. 4A and 5A). Interestingly, the PCA for CA mirrors the percent neutrophils concentration-response pattern – namely, 1 and 2 ppm-exposed samples cluster nearer to each other than to the FA-exposed samples and this threshold-like exposure-response pattern can largely be explained by PC1. The PCA for AM samples is more complex, with treatment effects reflected in both PC1 and PC2.

**Figure 4.**
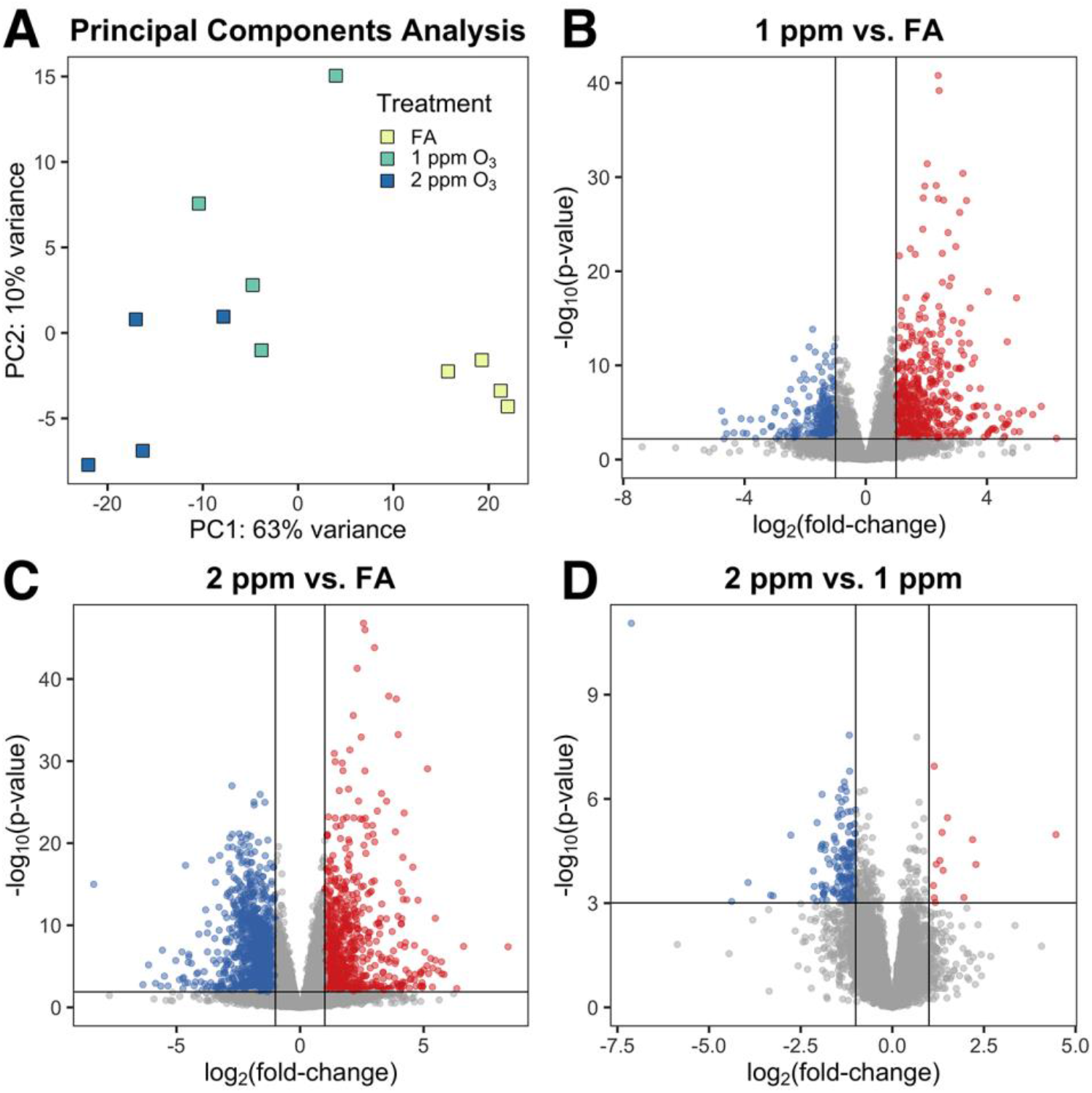
O_3_ exposure induces altered gene expression in conducting airways (CA). Gene expression of CA tissue from mice exposed to FA, 1 or 2 ppm O_3_. (A) Principal components analysis shows separation of the three treatment groups. Differential expression analysis revealed (B) 903 DEGs in 1 ppm versus FA airways, (C) 2,148 DEGs in 2 ppm versus FA airways, and (D) 188 DEGs in 2 ppm versus FA airways. (n=4 per treatment group, fold-change cutoff = 2, Benjamini-Hochberg adjusted p-value < 0.05)

**Figure 5.**
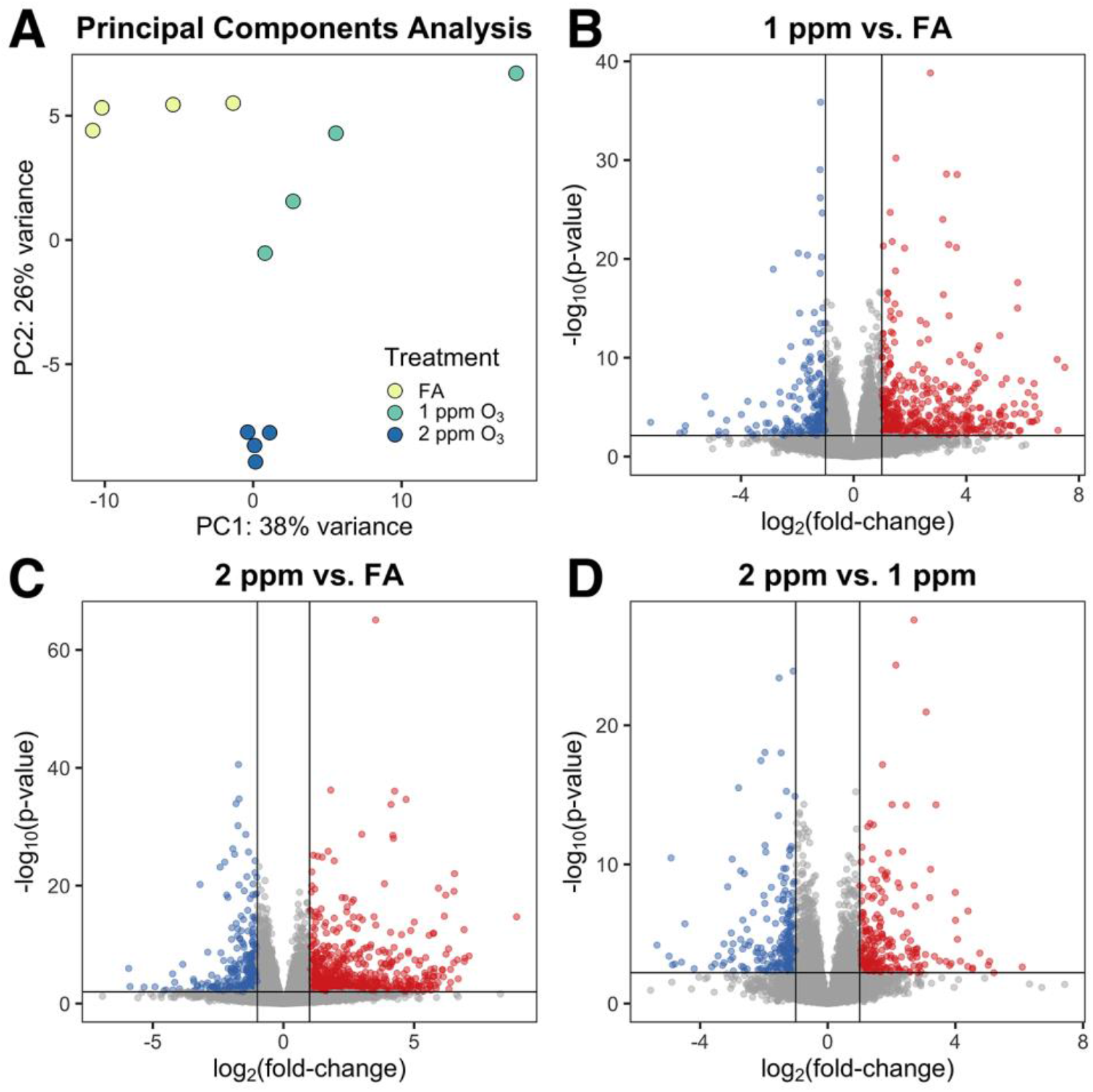
Airway macrophage (AM) gene expression is altered by O_3_ exposure. Gene expression in pooled AM from mice exposed to FA, 1 or 2 ppm O_3_. (A) Principal components analysis displays separation of AMs across all three treatment groups. Differential expression analysis revealed (B) 693 DEGs in 1 ppm versus FA AM samples, (C) 971 DEGs in 2 ppm versus FA AM samples, and (D) 467 DEGs in 2 versus 1 ppm AM samples. (n=4 pools per treatment group, fold-change cutoff = 2, Benjamini-Hochberg adjusted p-value < 0.05)

We identified differentially expressed genes (DEGs) by performing all pairwise comparisons between treatment groups within a tissue compartment. A summary of results from this analysis is included in Table 1, and all significantly DEGs (defined as FDR < 0.05 and absolute log_2_ fold-change > 1) for each treatment comparison and compartment are provided in Supplemental Material. In CA samples, we identified 903 DEGs between 1 ppm and FA, 2,148 DEGs between 2 ppm and FA, and 188 DEGs between 2 ppm and 1 ppm (Figure 4B-D). Generally, the range of fold-changes was greater within the 2 ppm versus FA comparison than within the 1 ppm versus FA comparison. In AM samples, we identified 694 DEGs between 1 ppm and FA, 972 DEGs between 2 ppm and FA, and 467 DEGs between 2 ppm and 1 ppm (Figure 5B-D). Similar to CA samples, the range of fold-changes was wider in the 2 ppm versus FA comparison than within the 1 ppm versus FA comparison in AM samples.

**Table 1.** The number of differentially expressed genes, by tissue compartment and treatment comparison. All genes had an absolute fold-change cutoff of 2 (i.e., absolute log_2_FC > 1).

| **Tissue** | **Comparison** | **FDR < 0.05** | **FDR < 0.1** |
| --- | --- | --- | --- |
| Conducting<br>airways | *1 ppm vs FA* | 903 | 1048 |
|  | *2 ppm vs FA* | 2148 | 2382 |
|  | *2 vs 1 ppm* | 188 | 255 |
| Airway<br>macrophages | *1 ppm vs FA* | 693 | 859 |
|  | *2 ppm vs FA* | 971 | 1216 |
|  | *2 vs 1 ppm* | 467 | 585 |

Many of the top genes with altered expression in the CA samples (Table 2) are those commonly associated with barrier function and epithelial organization (*Gjb3-5*, *Cldn2*/*4*, *Adam12, Tgm1*) (FRANK 2012; ESSELTINE AND LAIRD 2016), detoxification (*Mt1/2*, *Ugt1a6a/b*, *Gstm1/2/6*), and DNA replication (*Mcm5*-*7*, *Cdt1, Orc1*) (RIERA 2017). Notably, within CA samples, we observed an increase in expression of genes associated with proliferative responses (*Pcna*, *Cdh3*) concordant with our histopathological analysis. Furthermore, we noted a concentration-dependent decrease in expression of ciliated and club cell markers (*Foxj1*, *Cyp2f2*), corresponding with the loss of ciliated cells observed in the airways (HOGAN *et al*. 2014). While we observed no loss in protein expression of CCSP in our histopathological analysis, we did see a significant decrease in expression of *Scgb1a1* (the gene encoding CCSP) as well as *Scgb3a2*.

**Table 2.**
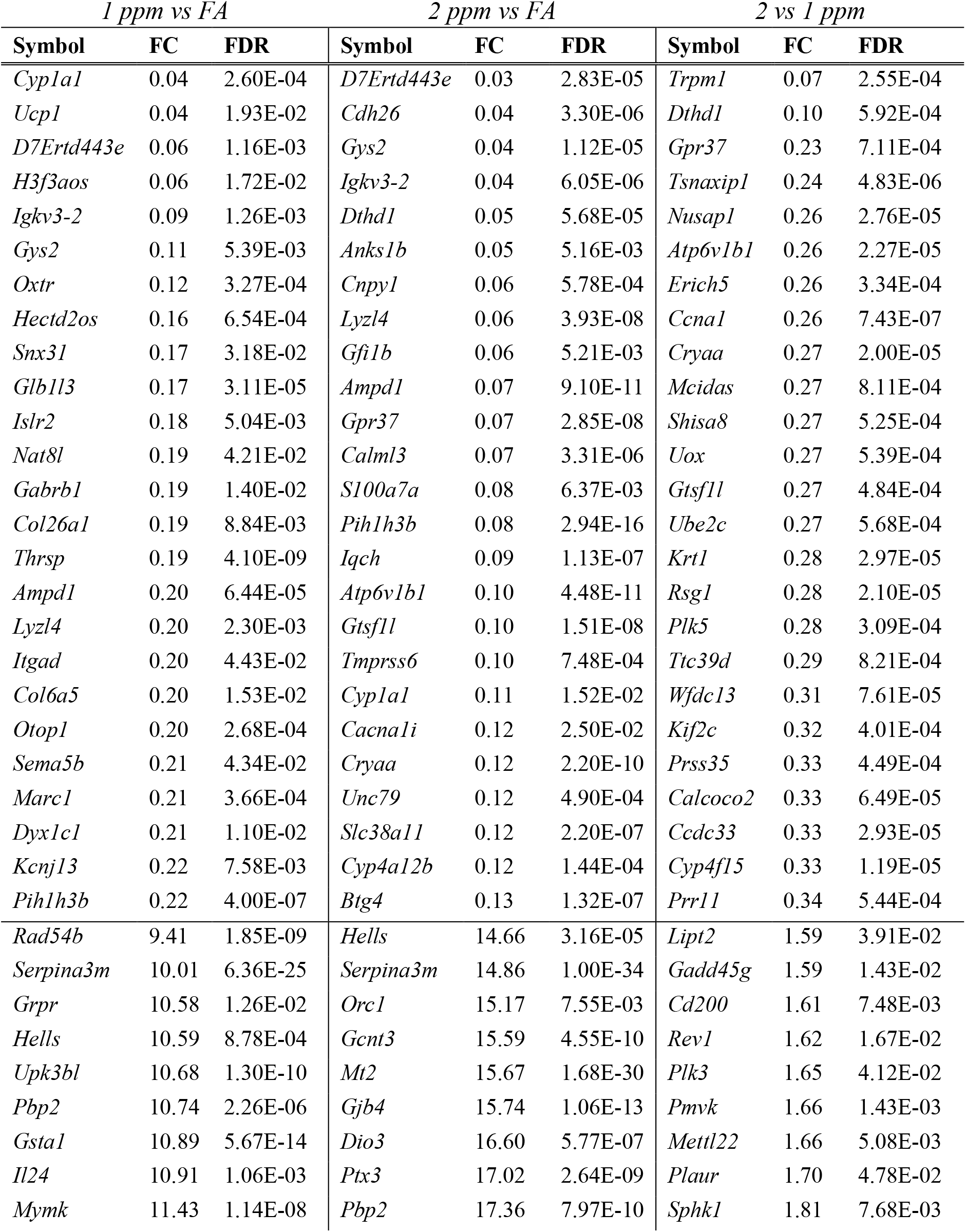
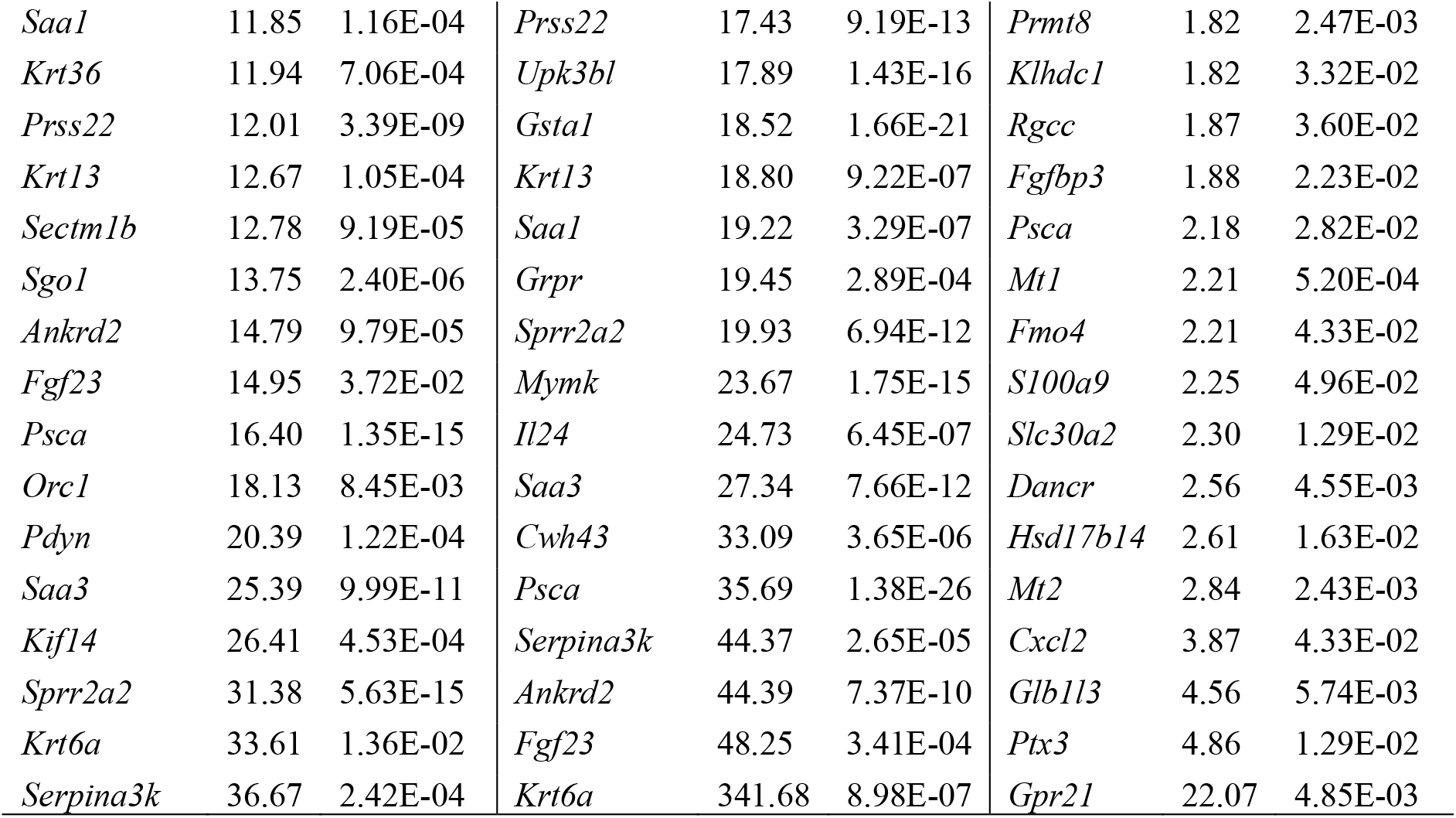
Top 50 most differentially expressed genes (25 downregulated, 25 upregulated) for each treatment comparison within conducting airways.

Within AM samples, genes involved in canonical immunological functions (*Ccl17*, *Slpi*, and *Ccl22*) and extracellular matrix organization and remodeling (*Krt7*, *Krt8*, *Krt18*) were highly differentially expressed (Table 3). Additionally, we observed differential expression of a number of genes that have been previously implicated in AM-mediated O_3_ responses including macrophage polarization genes (M1: *Cd80*, *Tnf*, *Ccl5*, *Tlr2*; M2: *Arg1*, *Retnla, Ccl24, Cxcl10*), proteases (*Mmp8*, *Mmp9, Mmp12, Ctsd, Ctse, Ctsf)*, and surfactant proteins (*S*ftpa1, *Sftpc*, *Sftpd*) (LASKIN *et al*. 2011; LASKIN *et al*. 2019).

To examine gene expression trends as a function of O_3_ concentration, we binned genes based on their expression pattern into four main groups, as illustrated in Figure 6A. We refer to the first group as monotonic, where genes showed a linear concentration-response pattern of expression. The second and third groups of genes exhibited threshold effects apparent at 1 or 2 ppm O_3_, respectively. Finally, the fourth group contained genes that were differentially expressed at 1 ppm versus both FA and 2 ppm O_3_. These four groups were further subdivided into eight distinct classes based on the directionality (up vs. down) of expression.

**Table 3.**
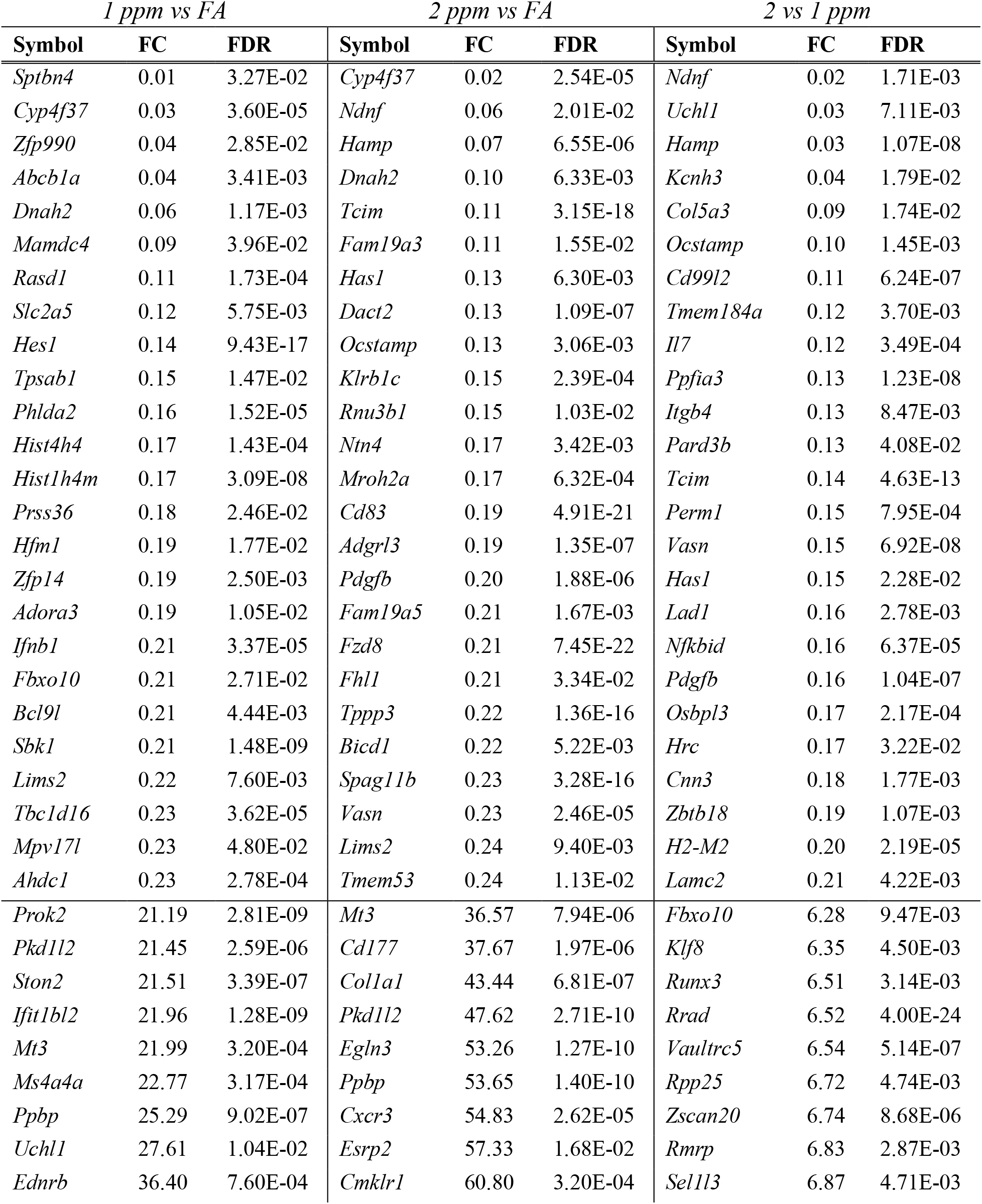
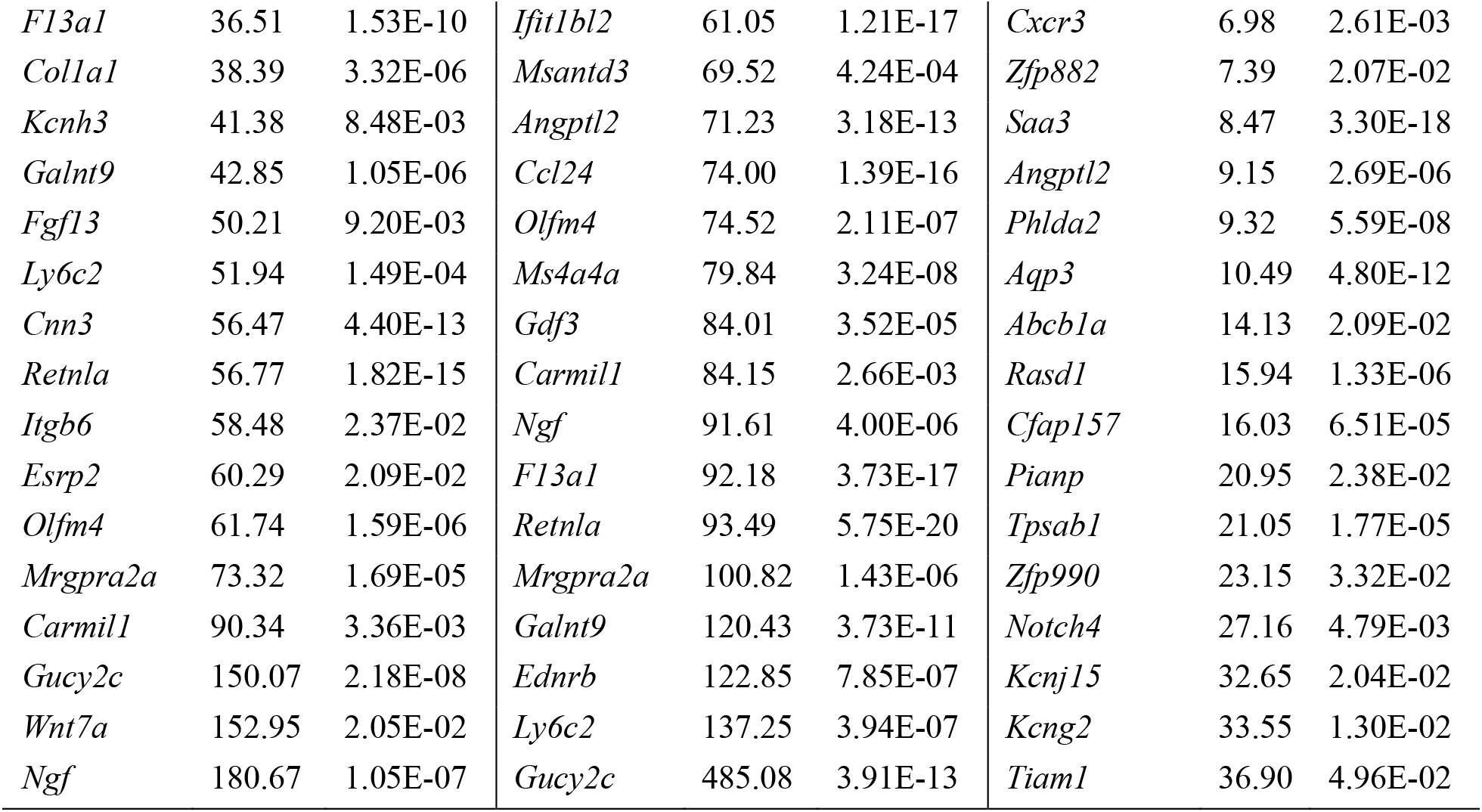
Top 50 most differentially expressed genes (25 downregulated, 25 upregulated) for each treatment comparison within airway macrophages.

**Figure 6.**
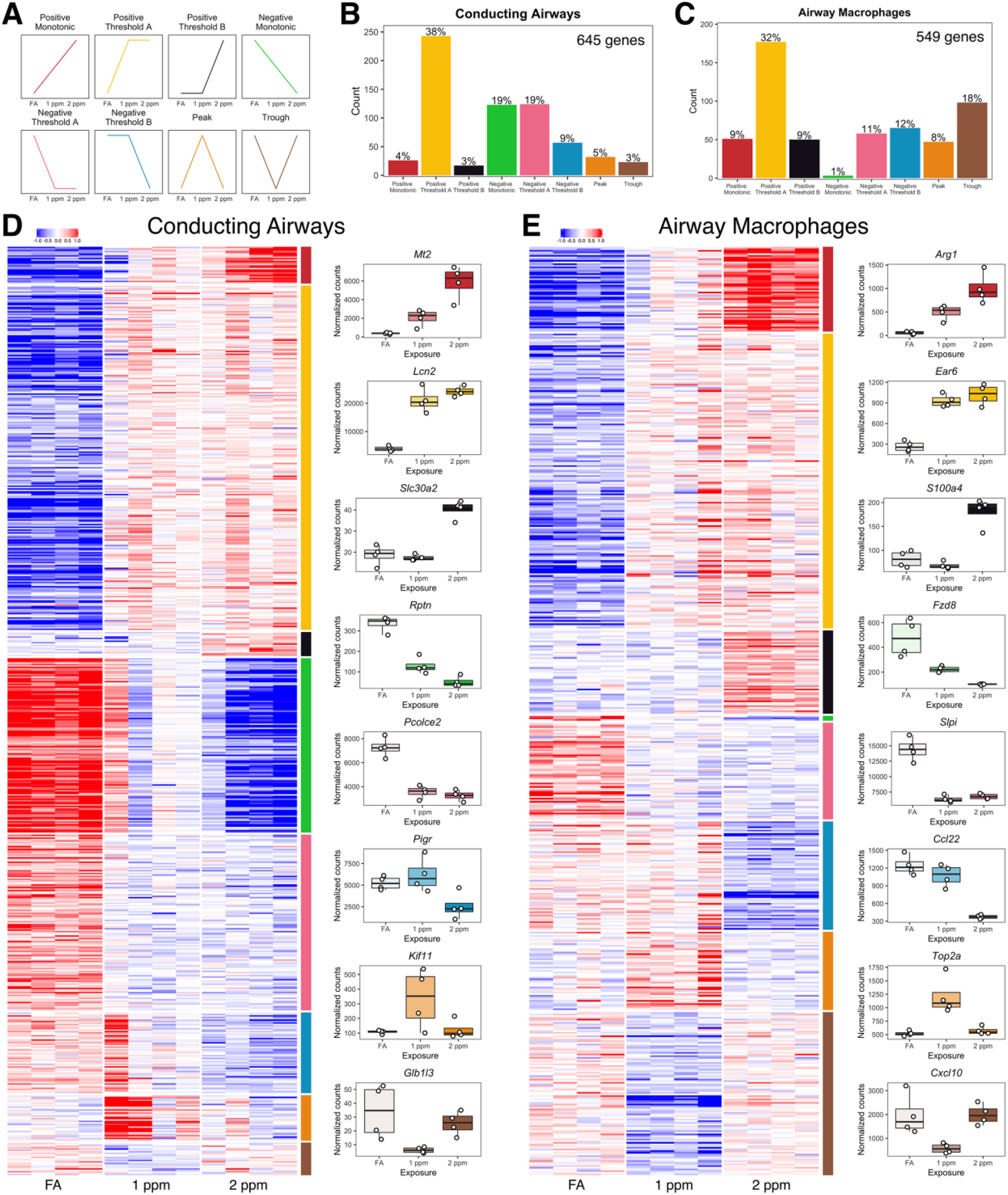
A subset of differentially expressed genes in both conducting airways and airway macrophages can be categorized into one of 8 distinct expression patterns. Schematic graphs of each category are displayed in panel A, and their order and the color assigned to each pattern is consistent throughout the figure. The frequency of genes binned into each trend category within (B) conducting airways and (C) airway macrophages. Heatmaps depict row-centered expression of categorized gene from (D) conducting airways and (E) airway macrophages, arranged by expression pattern. Boxplots of genes exemplifying each pattern are arranged to the right of each heatmap.

Overall, the number of genes that were categorized was similar between CA and AM (645 and 549, respectively; Figure 6B-C, Supplemental Table 2). There was a higher frequency of genes categorized as negative monotonic, negative threshold A (threshold across 1 and 2 ppm), and positive threshold A in CA versus in AM. Additionally, compared to CA, a larger number of AM genes were classified as positive monotonic, positive threshold B (threshold across FA and 1 ppm) and trough. After assigning DEGs to the 8 categories outlined above, we used the online Enrichr tool (KULESHOV *et al*. 2016) to explore functional annotations associated with each expression pattern (Supplemental Table 3). In CA, a number of pathways associated with protease function and inhibition were enriched within the positive monotonic category. Additionally, pathways involved in immunological signaling and defense responses were enriched in negative threshold B, and pathways regulating cell cycle progression and DNA damage responses were enriched in the peak category. For AM expression patterns, cytokine-cytokine receptor interactions and signaling gene sets were enriched in the positive monotonic category, DNA damage and break repair pathways were enriched in the positive threshold A category, and heat shock response pathways were enriched in the trough category.

We also compared DEGs across tissue compartments (Fig. 7, Supplemental Table 4). We identified 121 genes that were DE in both CA and AM comparing 1 ppm versus FA (Fig. 7A). For the 1 ppm versus FA contrast, only 13% of CA DEGs were shared with AM DEGs. Likewise, for the 2 ppm versus FA contrast, 8% of CA DEGs were shared with AM DEGs. Genes that were differentially expressed in both CA and AM after 1 ppm O_3_ included those involved in antioxidant responses (*Gsta1*), cell cycle progression and DNA replication (*Cdk1*, *Mcm2*, *Mcm6*, *Mcm8*, *Mcm10*), and acute phase proteins (*Clu*, *Lcn2*, *Saa3*). After 2 ppm O_3_ exposure, CA and AM exhibited differential expression of genes involved in immune signaling (*Ccl17*, *Slpi*, *Cx3cl1*) in addition to some of the cell cycle-associated and acute phase genes (*Mcm2*, *Mcm10*, *Lcn2*, *Saa3*) that were differentially expressed after 1 ppm O_3_ exposure. Lastly, only one gene (*Hsph1*), a heat-shock protein, was differentially expressed in both CA and AM when comparing 2 ppm versus 1 ppm O_3_, albeit discordantly (up in CA, down in AM).

**Figure 7.**
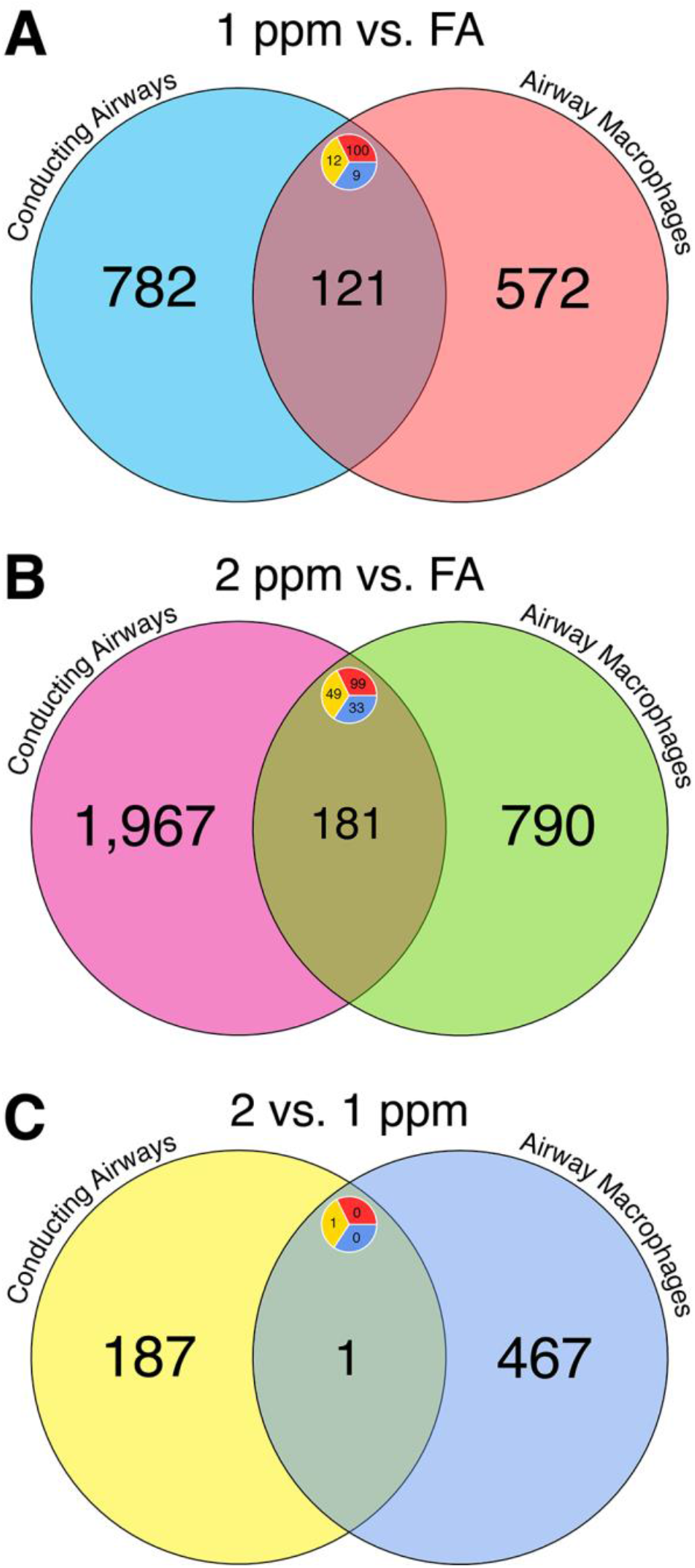
Comparison of differential gene expression results shows shared and unique responses to O_3_ across tissue compartments. Venn diagrams comparing lists of differentially expressed (DE) genes in conducting airway tissue and airway macrophages from (A) the 1 ppm versus FA comparison, (B) the 2 ppm versus FA comparison, and (C) the 2 versus 1 ppm comparison. Labels display the number of genes within a set. Inset circles display the number of upregulated (red), down regulated (blue), or genes with contrasting expression patterns (yellow) within a union set.

Finally, we compared our lists of DEGs to those published in three previous studies (GOHIL *et al*. 2003; KOOTER *et al*. 2007; GABEHART *et al*. 2014) (Supplemental Tables 5 & 6) of whole lung tissue to identify which genes are consistently differentially expressed, and to determine which DEGs from CA or AM are captured by whole lung transcriptomics. Notably, all comparisons had significant overlap between our lists of DEGs and those from previous studies, with CA samples overlapping with previous studies with a higher level of significance. The range of fold-enrichment represented by CA samples ranged from ∼3-13, while the range in AM samples was from ∼2.5-5 (Supplemental Table 7). We also used the vote-counting method of meta-analysis to identify genes that were consistently differentially expressed across multiple studies (Supplemental Table 8). In CA samples, 76 unique genes overlapped across all studies with our list of DEGs, with 5 genes represented in 2 studies and our own (*Cdk1*, *Lcn2*, *Mt1*, *Saa3*, *Serpina3n*). In AM samples, 38 unique genes overlapped with our DEGs, with 4 genes included in 2 other studies and our own (*Cdk1*, *Lcn2*, *S100a9*, *Saa3*). Finally, we compared the lists of overlapping genes between CA and AM samples to examine whether whole-lung transcriptomic studies captured compartment-specific gene expression patterns (Supplemental Table 9). Given that the proportion of CA-derived cells in the whole lung is greater than the proportion of AMs, we expected that previous studies would detect gene expression changes primarily reflective of CA. Indeed, 57 genes of the published whole lung DEGs overlapped with CA DEGs, while 19 genes were uniquely shared with AM DEGs (Supplemental Table 8).

### Pathway level effects of O_3_ exposure

In order to assess pathway level changes due to O_3_ in CA or AM, we performed Gene Set Variation Analysis (GSVA), which has been shown to be more powerful at detecting subtle changes in pathway level expression compared to previous methods (HÄNZELMANN 2013). This is because in GSVA, for each sample, the expression scores of a predefined set of genes (i.e., pathways) are summed and then differential expression analysis is performed on these aggregate scores. Thus, modest differences in the expression of genes that may not have reached statistical significance (for each gene individually) can still cumulatively result in differential expression of a gene set. The results of these analyses are summarized in Table 4, and all significantly enriched gene sets are reported in Supplemental Material. Given the exploratory nature of GSVA, we used a less stringent FDR of 0.1 for these analyses. In CA, 1,318 gene sets were differentially enriched in the 1 ppm versus FA comparison, and 2,377 gene sets were differentially enriched in the 2 ppm versus FA comparison. Of note, no gene sets were differentially enriched in the 2 versus 1 ppm comparison within CA. In AM samples, 1,514 gene sets were differentially enriched between 1 ppm versus FA, 2,931 gene sets between 2 ppm versus FA, and 1,223 gene sets between 2 versus 1 ppm.

**Table 4.**
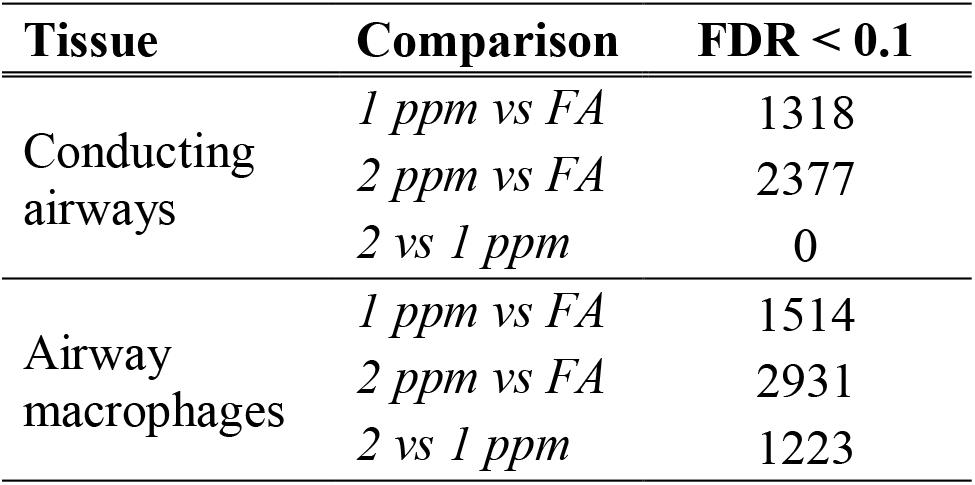
The number of differentially enriched gene sets, by tissue compartment and treatment comparison.

| **Tissue** | **Comparison** | **FDR < 0.1** |
| --- | --- | --- |
| Conducting<br>airways | *1 ppm vs FA* | 1318 |
|  | *2 ppm vs FA* | 2377 |
|  | *2 vs 1 ppm* | 0 |
| Airway<br>macrophages | *1 ppm vs FA* | 1514 |
|  | *2 ppm vs FA* | 2931 |
|  | *2 vs 1 ppm* | 1223 |

**Table 5.**
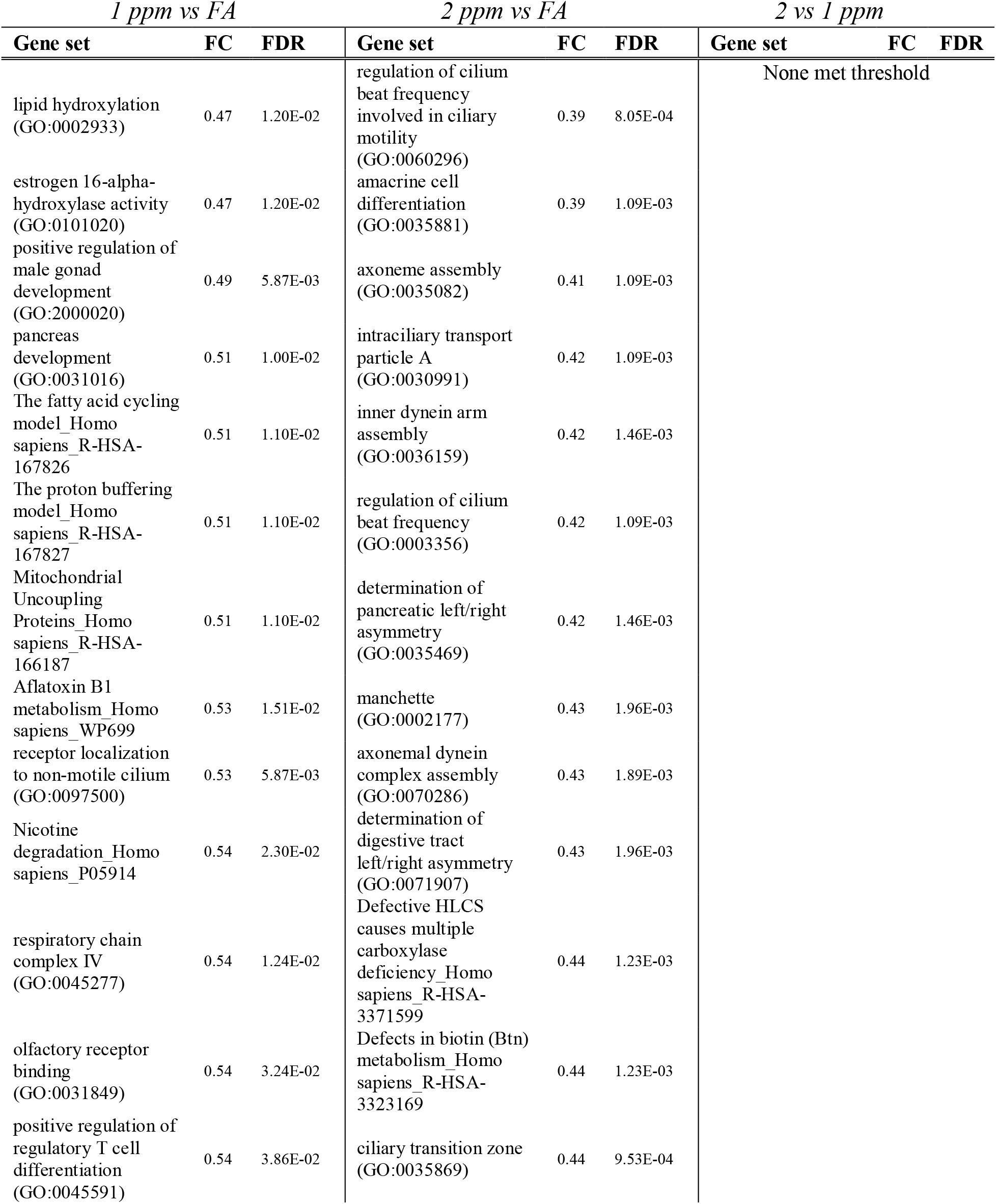
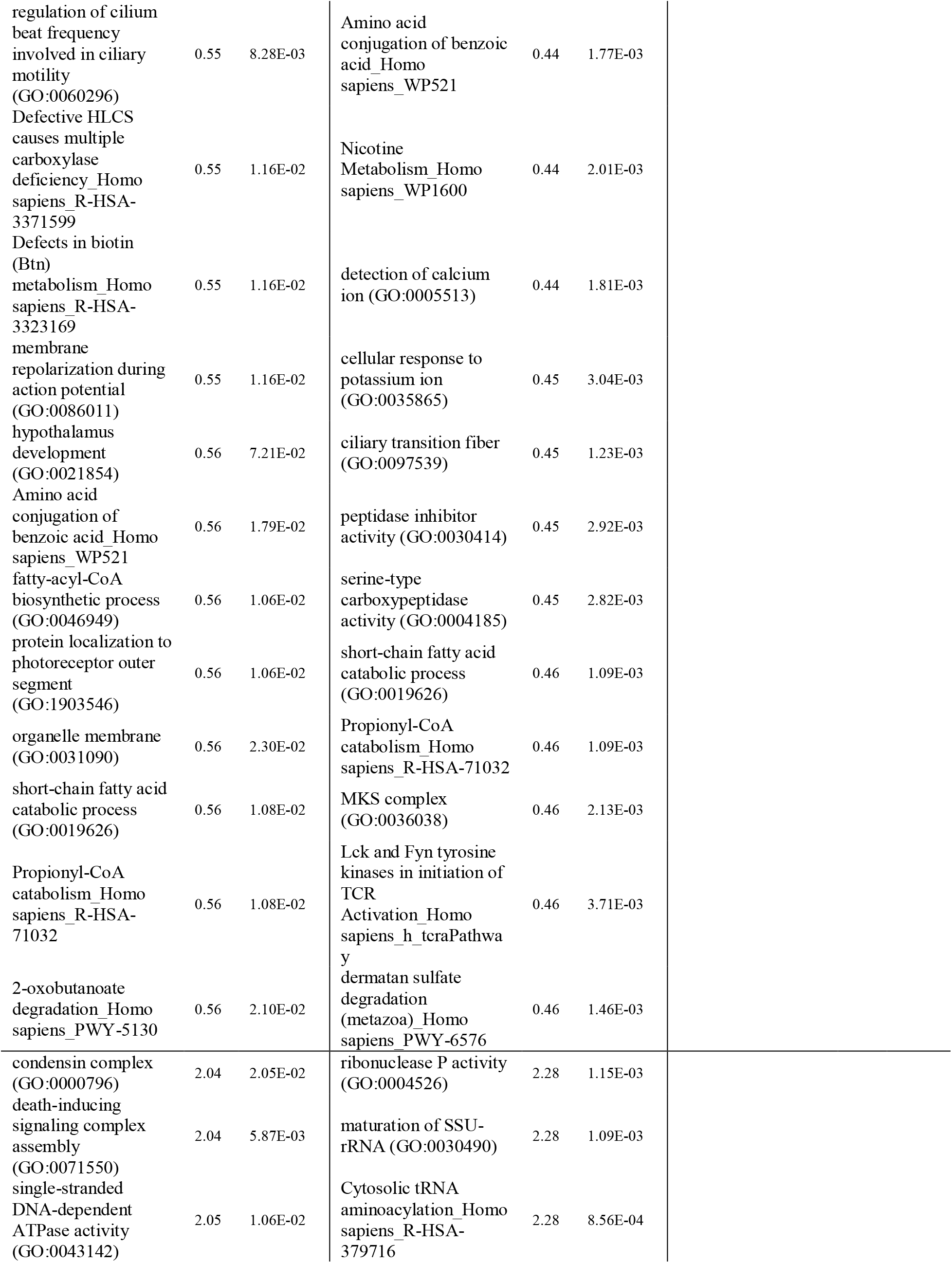
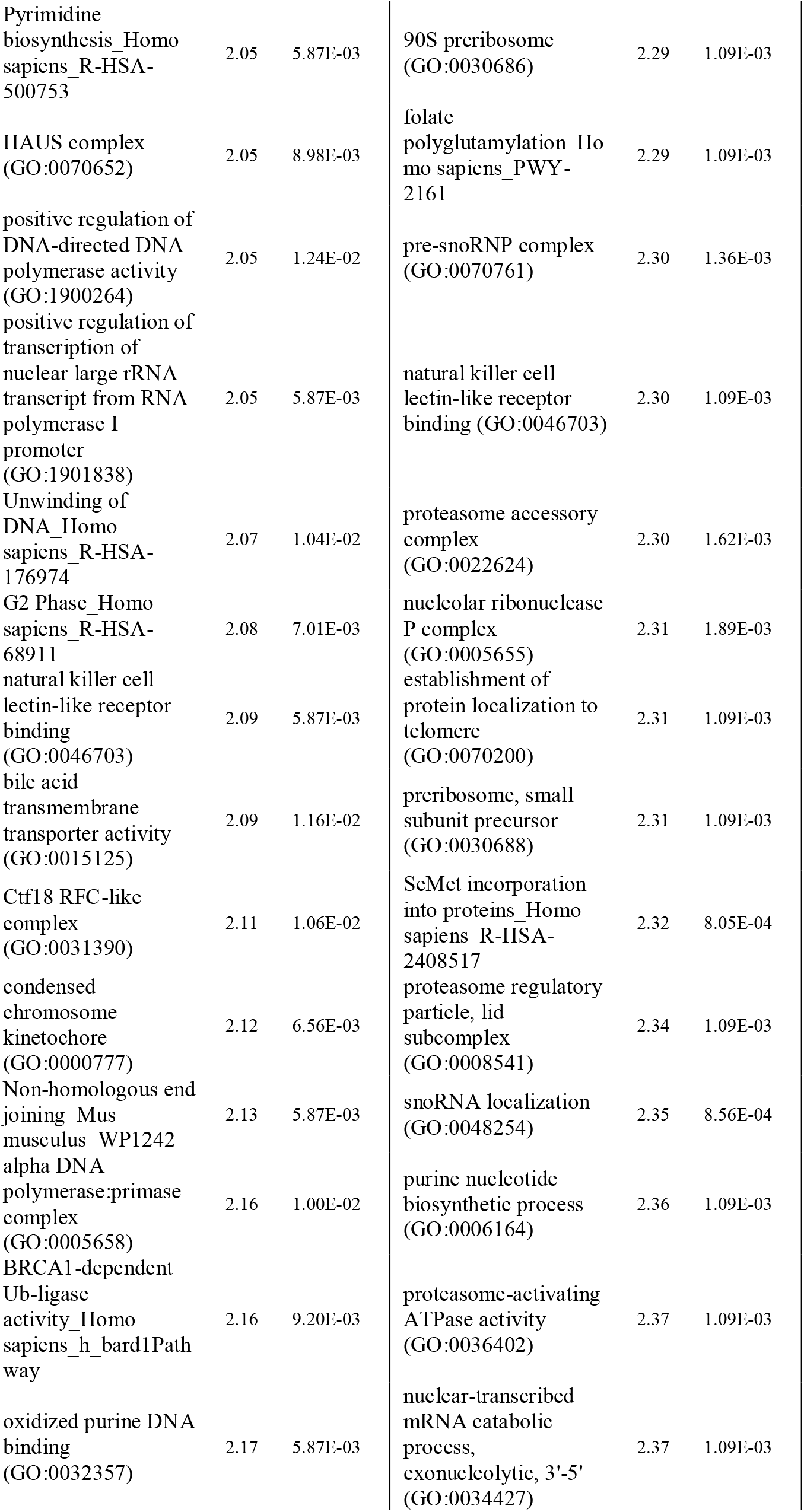
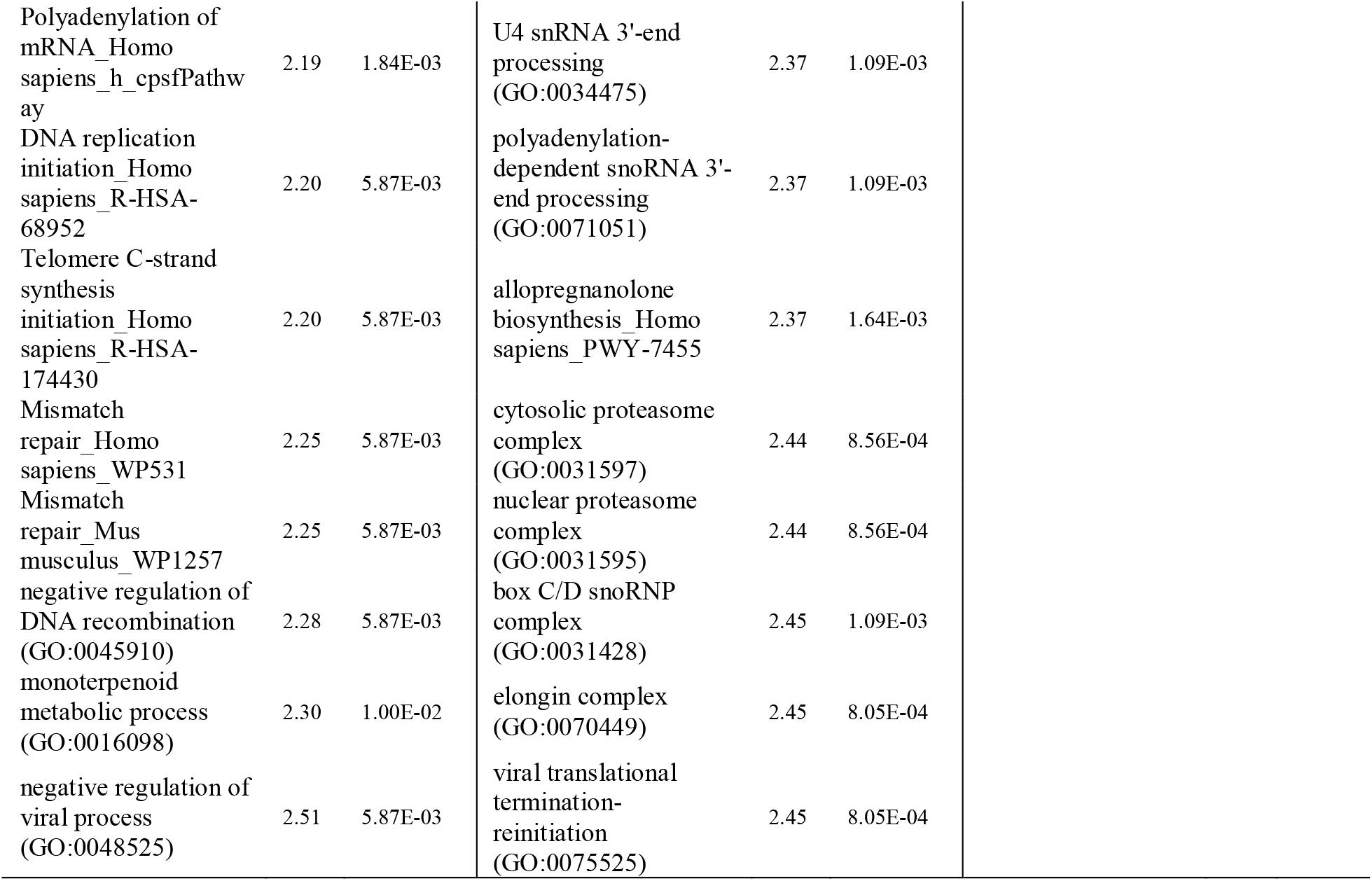
Top 50 most differentially enriched gene sets (25 down, 25 up) for each treatment comparison within conducting airways.

| *1 ppm vs FA* |  |  | *2 ppm vs FA* |  |  | *2 vs 1 ppm* |  |  |
| --- | --- | --- | --- | --- | --- | --- | --- | --- |
| Gene set | FC | FDR | Gene set | FC | FDR | Gene set | FC | FDR |
| lipid hydroxylation<br>(GO:0002933) | 0.47 | 1.20E-02 | regulation of cilium<br>beat frequency<br>involved in ciliary<br>motility<br>(GO:0060296) | 0.39 | 8.05E-04 | None met threshold |  |  |
| estrogen 16-alpha-<br>hydroxylase activity<br>(GO:0101020) | 0.47 | 1.20E-02 | amacrine cell<br>differentiation<br>(GO:0035881) | 0.39 | 1.09E-03 |  |  |  |
| positive regulation of<br>male gonad<br>development<br>(GO:2000020) | 0.49 | 5.87E-03 | axoneme assembly<br>(GO:0035082) | 0.41 | 1.09E-03 |  |  |  |
| pancreas<br>development<br>(GO:0031016) | 0.51 | 1.00E-02 | intraciliary transport<br>particle A<br>(GO:0030991) | 0.42 | 1.09E-03 |  |  |  |
| The fatty acid cycling<br>model_Homo<br>sapiens_R-HSA-167826 | 0.51 | 1.10E-02 | inner dynein arm<br>assembly<br>(GO:0036159) | 0.42 | 1.46E-03 |  |  |  |
| The proton buffering<br>model_Homo<br>sapiens_R-HSA-167827 | 0.51 | 1.10E-02 | regulation of cilium<br>beat frequency<br>(GO:0003356) | 0.42 | 1.09E-03 |  |  |  |
| Mitochondrial<br>Uncoupling<br>Proteins_Homo<br>sapiens_R-HSA-166187 | 0.51 | 1.10E-02 | determination of<br>pancreatic left/right<br>asymmetry<br>(GO:0035469) | 0.42 | 1.46E-03 |  |  |  |
| Aflatoxin B1<br>metabolism_Homo<br>sapiens_WP699 | 0.53 | 1.51E-02 | manchette<br>(GO:0002177) | 0.43 | 1.96E-03 |  |  |  |
| receptor localization<br>to non-motile cilium<br>(GO:0097500) | 0.53 | 5.87E-03 | axonemal dynein<br>complex assembly<br>(GO:0070286) | 0.43 | 1.89E-03 |  |  |  |
| Nicotine<br>degradation_Homo<br>sapiens_P05914 | 0.54 | 2.30E-02 | determination of<br>digestive tract<br>left/right asymmetry<br>(GO:0071907) | 0.43 | 1.96E-03 |  |  |  |
| respiratory chain<br>complex IV<br>(GO:0045277) | 0.54 | 1.24E-02 | Defective HLCS<br>causes multiple<br>carboxylase<br>deficiency_Homo<br>sapiens_R-HSA-3371599 | 0.44 | 1.23E-03 |  |  |  |
| olfactory receptor<br>binding<br>(GO:0031849) | 0.54 | 3.24E-02 | Defects in biotin (Btn)<br>metabolism_Homo<br>sapiens_R-HSA-3323169 | 0.44 | 1.23E-03 |  |  |  |
| positive regulation of<br>regulatory T cell<br>differentiation<br>(GO:0045591) | 0.54 | 3.86E-02 | ciliary transition zone<br>(GO:0035869) | 0.44 | 9.53E-04 |  |  |  |

|  |  |  |  |  |  |
| --- | --- | --- | --- | --- | --- |
| regulation of cilium beat frequency involved in ciliary motility (GO:0060296) | 0.55 | 8.28E-03 | Amino acid conjugation of benzoic acid_Homo sapiens_WP521 | 0.44 | 1.77E-03 |
| Defective HLCS causes multiple carboxylase deficiency_Homo sapiens_R-HSA-3371599 | 0.55 | 1.16E-02 | Nicotine Metabolism_Homo sapiens_WP1600 | 0.44 | 2.01E-03 |
| Defects in biotin (Btn) metabolism_Homo sapiens_R-HSA-3323169 | 0.55 | 1.16E-02 | detection of calcium ion (GO:0005513) | 0.44 | 1.81E-03 |
| membrane repolarization during action potential (GO:0086011) | 0.55 | 1.16E-02 | cellular response to potassium ion (GO:0035865) | 0.45 | 3.04E-03 |
| hypothalamus development (GO:0021854) | 0.56 | 7.21E-02 | ciliary transition fiber (GO:0097539) | 0.45 | 1.23E-03 |
| Amino acid conjugation of benzoic acid_Homo sapiens_WP521 | 0.56 | 1.79E-02 | peptidase inhibitor activity (GO:0030414) | 0.45 | 2.92E-03 |
| fatty-acyl-CoA biosynthetic process (GO:0046949) | 0.56 | 1.06E-02 | serine-type carboxypeptidase activity (GO:0004185) | 0.45 | 2.82E-03 |
| protein localization to photoreceptor outer segment (GO:1903546) | 0.56 | 1.06E-02 | short-chain fatty acid catabolic process (GO:0019626) | 0.46 | 1.09E-03 |
| organelle membrane (GO:0031090) | 0.56 | 2.30E-02 | Propionyl-CoA catabolism_Homo sapiens_R-HSA-71032 | 0.46 | 1.09E-03 |
| short-chain fatty acid catabolic process (GO:0019626) | 0.56 | 1.08E-02 | MKS complex (GO:0036038) | 0.46 | 2.13E-03 |
| Propionyl-CoA catabolism_Homo sapiens_R-HSA-71032 | 0.56 | 1.08E-02 | Lck and Fyn tyrosine kinases in initiation of TCR Activation_Homo sapiens_h_tcraPathway | 0.46 | 3.71E-03 |
| 2-oxobutanoate degradation_Homo sapiens_PWY-5130 | 0.56 | 2.10E-02 | dermatan sulfate degradation (metazoa)_Homo sapiens_PWY-6576 | 0.46 | 1.46E-03 |
| condensin complex (GO:0000796) | 2.04 | 2.05E-02 | ribonuclease P activity (GO:0004526) | 2.28 | 1.15E-03 |
| death-inducing signaling complex assembly (GO:0071550) | 2.04 | 5.87E-03 | maturation of SSU-rRNA (GO:0030490) | 2.28 | 1.09E-03 |
| single-stranded DNA-dependent ATPase activity (GO:0043142) | 2.05 | 1.06E-02 | Cytosolic tRNA aminoacylation_Homo sapiens_R-HSA-379716 | 2.28 | 8.56E-04 |
| Pyrimidine biosynthesis_Homo sapiens_R-HSA-500753 | 2.05 | 5.87E-03 | 90S preribosome (GO:0030686) | 2.29 | 1.09E-03 |
| HAUS complex (GO:0070652) | 2.05 | 8.98E-03 | folate polyglutamylation_Homo sapiens_PWY-2161 | 2.29 | 1.09E-03 |
| positive regulation of DNA-directed DNA polymerase activity (GO:1900264) | 2.05 | 1.24E-02 | pre-snoRNP complex (GO:0070761) | 2.30 | 1.36E-03 |
| positive regulation of transcription of nuclear large rRNA transcript from RNA polymerase I promoter (GO:1901838) | 2.05 | 5.87E-03 | natural killer cell lectin-like receptor binding (GO:0046703) | 2.30 | 1.09E-03 |
| Unwinding of DNA_Homo sapiens_R-HSA-176974 | 2.07 | 1.04E-02 | proteasome accessory complex (GO:0022624) | 2.30 | 1.62E-03 |
| G2 Phase_Homo sapiens_R-HSA-68911 | 2.08 | 7.01E-03 | nucleolar ribonuclease P complex (GO:0005655) | 2.31 | 1.89E-03 |
| natural killer cell lectin-like receptor binding (GO:0046703) | 2.09 | 5.87E-03 | establishment of protein localization to telomere (GO:0070200) | 2.31 | 1.09E-03 |
| bile acid transmembrane transporter activity (GO:0015125) | 2.09 | 1.16E-02 | preribosome, small subunit precursor (GO:0030688) | 2.31 | 1.09E-03 |
| Ctf18 RFC-like complex (GO:0031390) | 2.11 | 1.06E-02 | SeMet incorporation into proteins_Homo sapiens_R-HSA-2408517 | 2.32 | 8.05E-04 |
| condensed chromosome kinetochore (GO:0000777) | 2.12 | 6.56E-03 | proteasome regulatory particle, lid subcomplex (GO:0008541) | 2.34 | 1.09E-03 |
| Non-homologous end joining_Mus musculus_WP1242 | 2.13 | 5.87E-03 | snoRNA localization (GO:0048254) | 2.35 | 8.56E-04 |
| alpha DNA polymerase:primase complex (GO:0005658) | 2.16 | 1.00E-02 | purine nucleotide biosynthetic process (GO:0006164) | 2.36 | 1.09E-03 |
| BRCA1-dependent Ub-ligase activity_Homo sapiens_h_bard1Pathway | 2.16 | 9.20E-03 | proteasome-activating ATPase activity (GO:0036402) | 2.37 | 1.09E-03 |
| oxidized purine DNA binding (GO:0032357) | 2.17 | 5.87E-03 | nuclear-transcribed mRNA catabolic process, exonucleolytic, 3'-5' (GO:0034427) | 2.37 | 1.09E-03 |
| Polyadenylation of mRNA_Homo sapiens_h_cpsfPathway | 2.19 | 1.84E-03 | U4 snRNA 3'-end processing (GO:0034475) | 2.37 | 1.09E-03 |
| DNA replication initiation_Homo sapiens_R-HSA-68952 | 2.20 | 5.87E-03 | polyadenylation-dependent snoRNA 3'-end processing (GO:0071051) | 2.37 | 1.09E-03 |
| Telomere C-strand synthesis initiation_Homo sapiens_R-HSA-174430 | 2.20 | 5.87E-03 | allopregnanolone biosynthesis_Homo sapiens_PWY-7455 | 2.37 | 1.64E-03 |
| Mismatch repair_Homo sapiens_WP531 | 2.25 | 5.87E-03 | cytosolic proteasome complex (GO:0031597) | 2.44 | 8.56E-04 |
| Mismatch repair_Mus musculus_WP1257 | 2.25 | 5.87E-03 | nuclear proteasome complex (GO:0031595) | 2.44 | 8.56E-04 |
| negative regulation of DNA recombination (GO:0045910) | 2.28 | 5.87E-03 | box C/D snoRNP complex (GO:0031428) | 2.45 | 1.09E-03 |
| monoterpenoid metabolic process (GO:0016098) | 2.30 | 1.00E-02 | elongin complex (GO:0070449) | 2.45 | 8.05E-04 |
| negative regulation of viral process (GO:0048525) | 2.51 | 5.87E-03 | viral translational termination-reinitiation (GO:0075525) | 2.45 | 8.05E-04 |

**Table 6.**
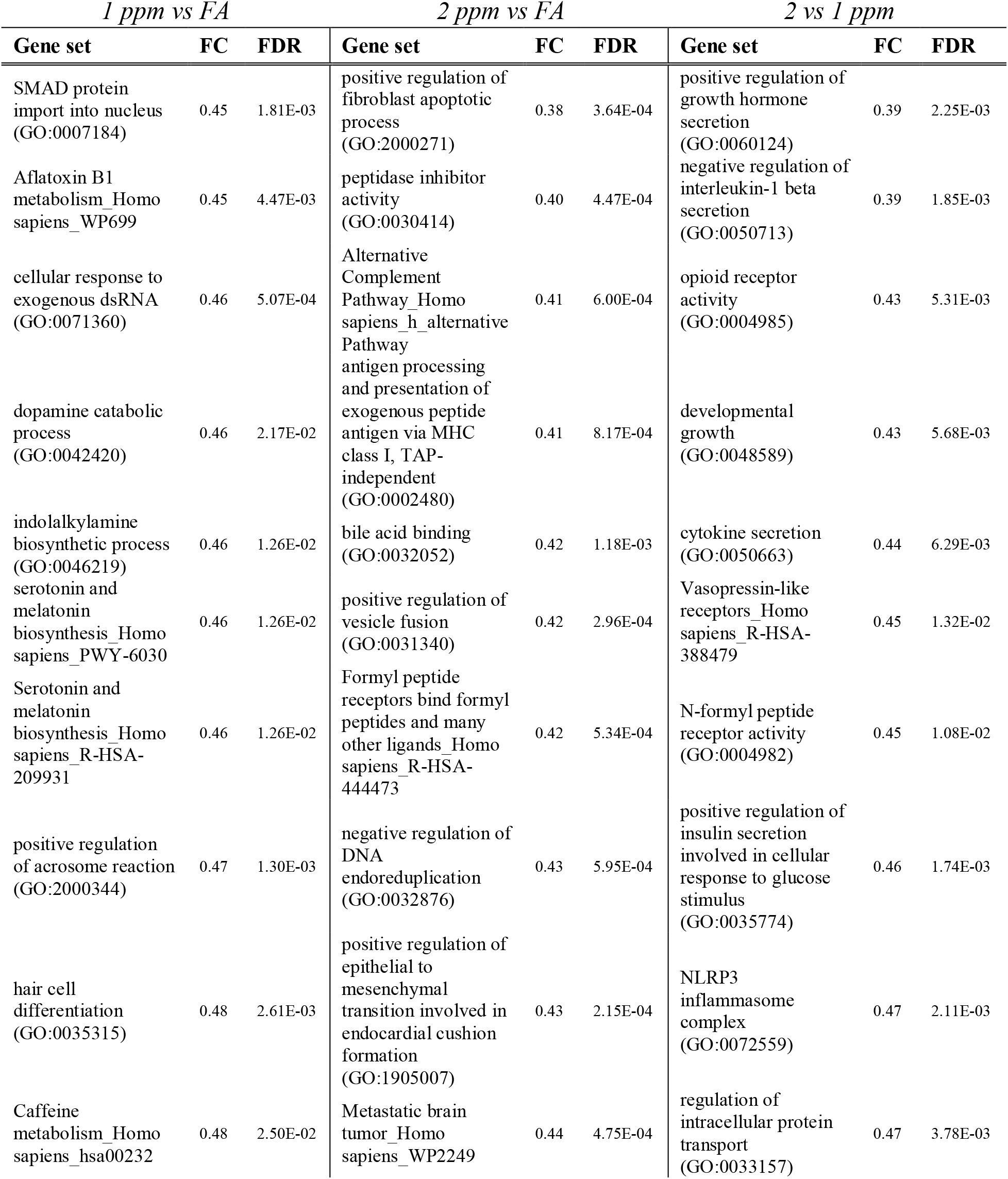
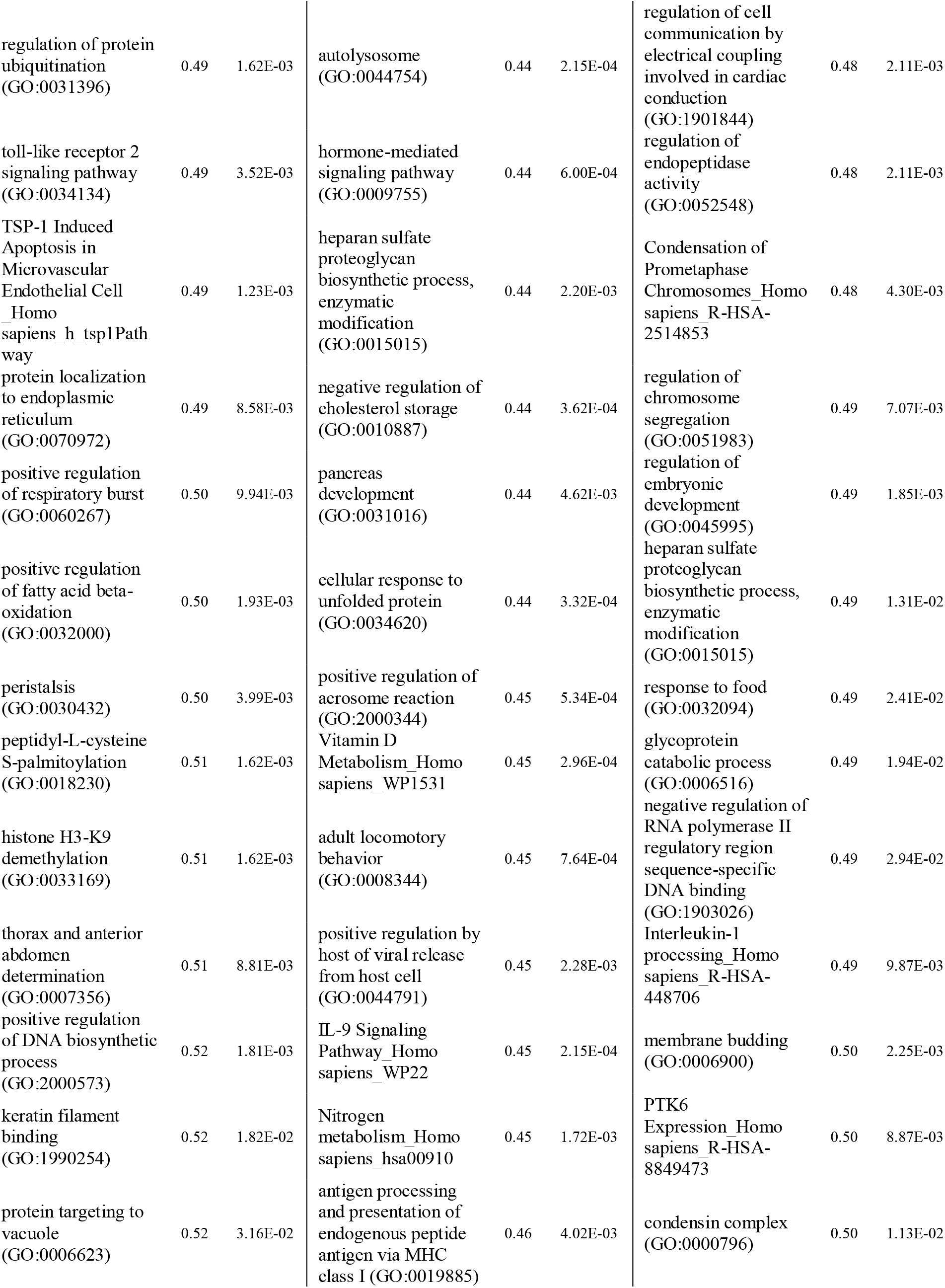
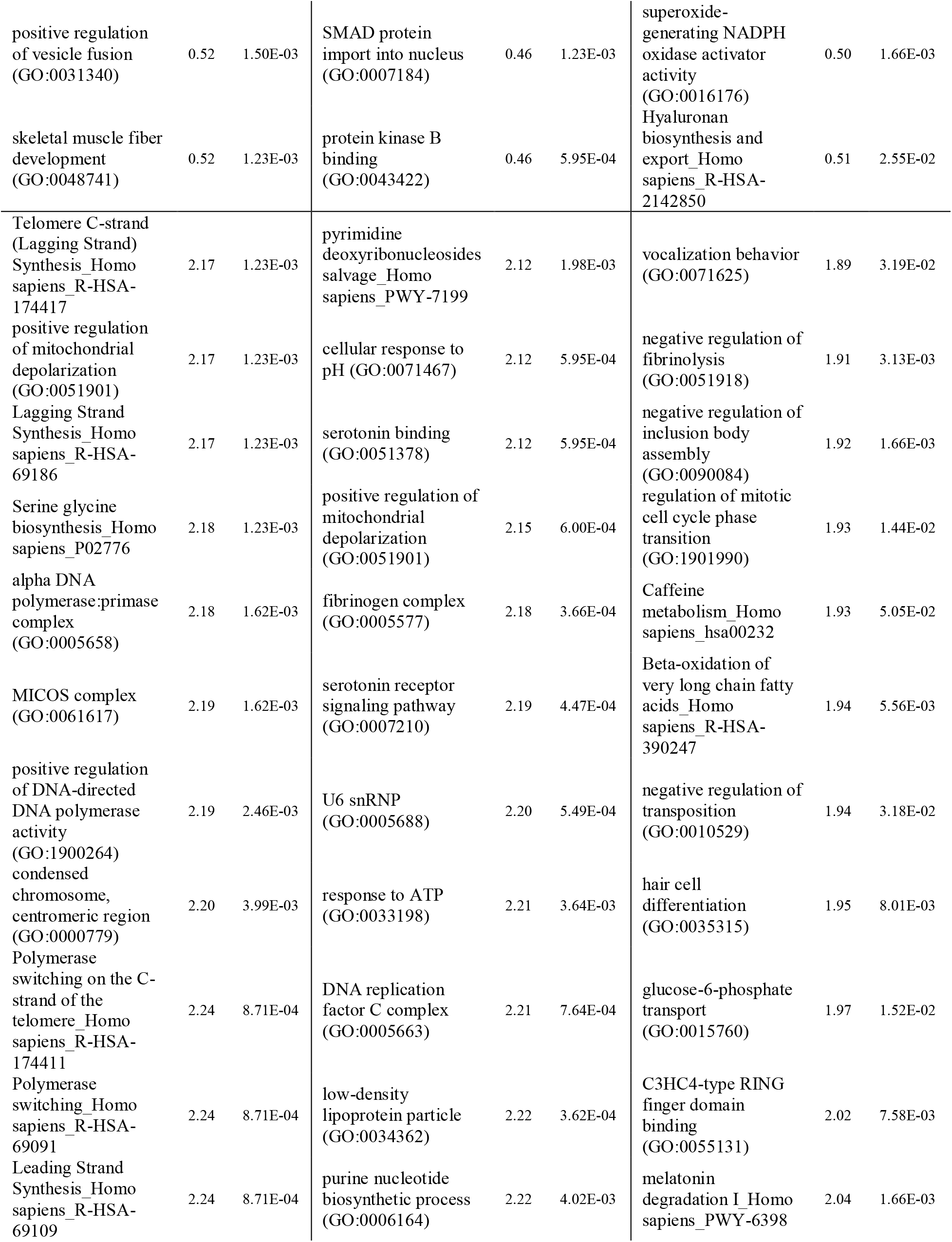
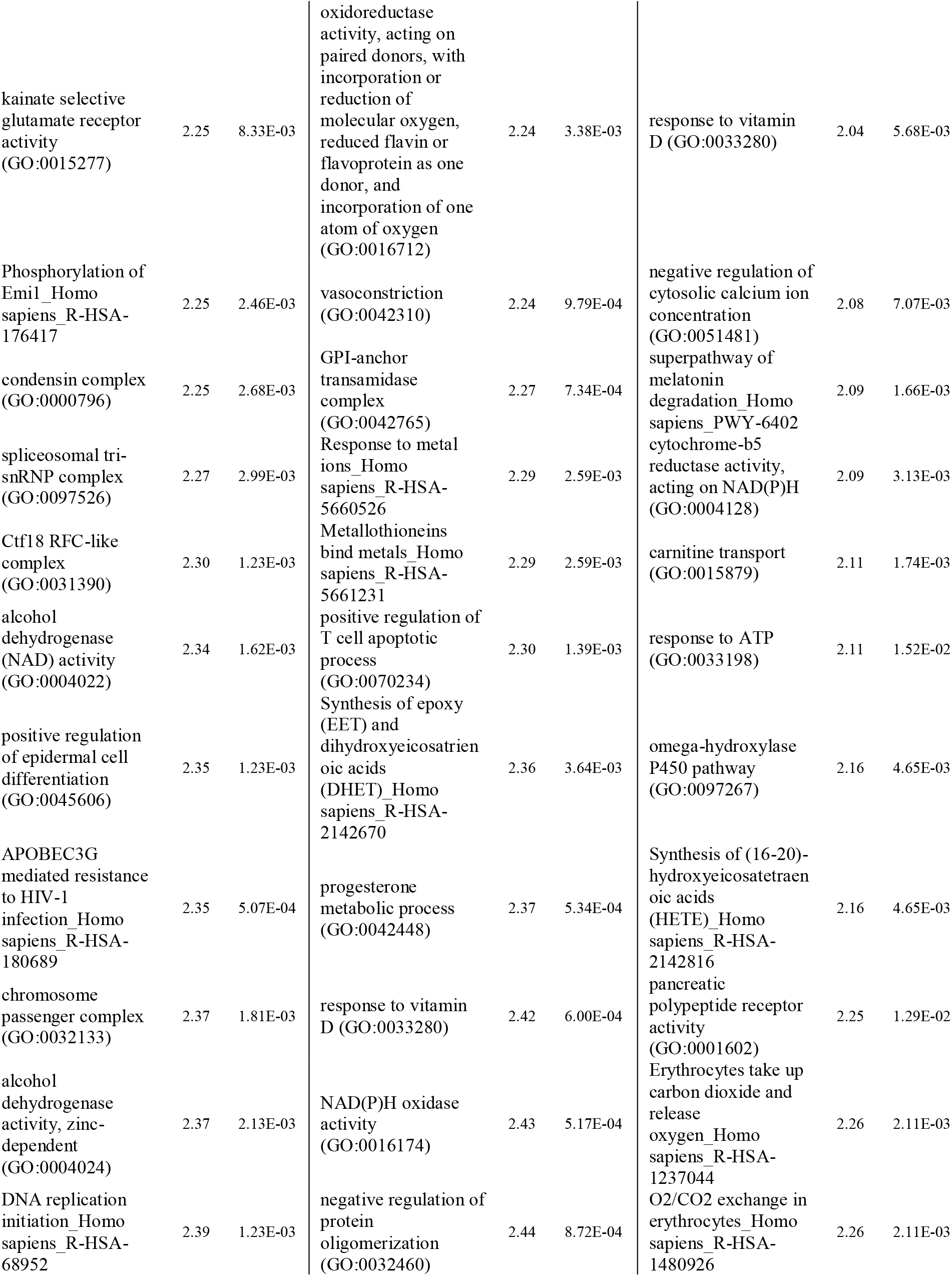
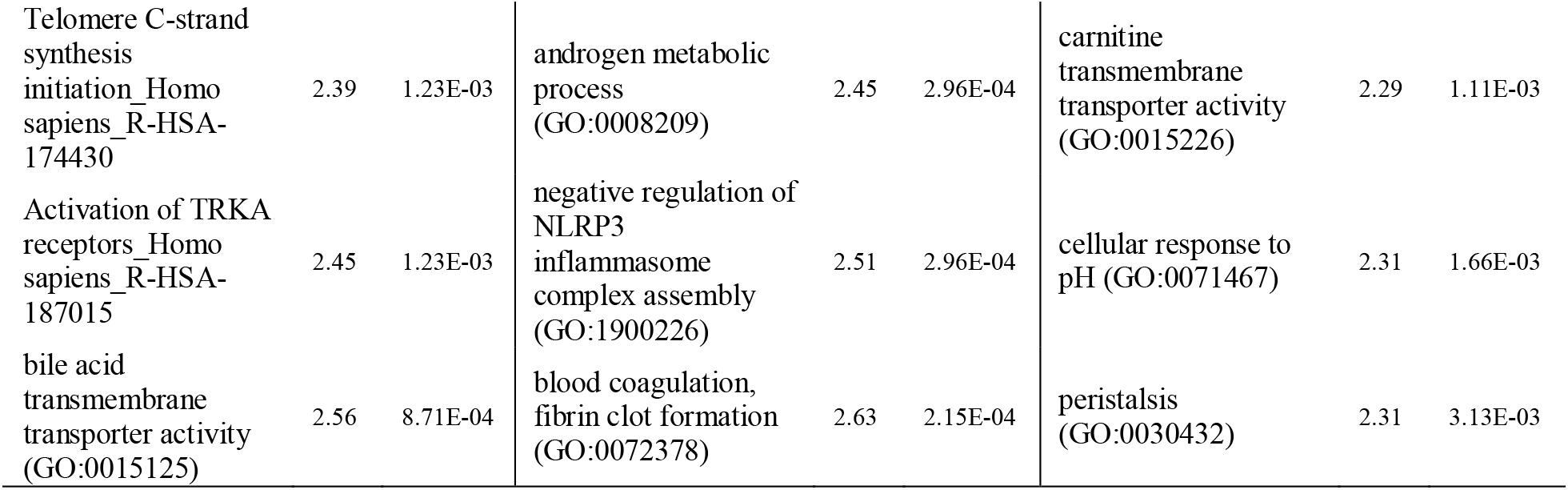
Top 50 most differentially enriched gene sets (25 down, 25 up) for each treatment comparison within airway macrophages.

We noted a clear decrease in expression of gene sets related to cilia function in CA, in line with the epithelial injury and ciliated cell loss in our histological analysis. In contrast, there were multiple gene sets related to DNA replication and DNA damage repair that were increased in response to 1 ppm (versus FA), and pathways involved in proteasome function and activity were upregulated in response to 2 ppm O_3_.

Within AM, many pathways involved in oxidative stress and detoxification responses including metallothionein and alcohol dehydrogenase activity were increased due to 1 and 2 ppm O_3_. Fc-gamma-regulated phagocytosis, positive regulation of vesicle fusion and other endocytic pathways were down regulated after 1 and 2 ppm O_3_, giving support to previous reports that O_3_ disrupts AM phagocytosis (GILMOUR *et al*. 1991; JAKAB *et al*. 1995). Antigen processing and presentation by MHC-I and other related pathways were also decreased after 2 ppm O_3_. Pathways that were upregulated in AM after 2 versus 1 ppm include erythrocyte O_2_/CO_2_ exchange and arachidonic acid metabolism, while cellular communication pathways including secretion of various soluble mediators (e.g., growth factors, cytokines, and hormones) were downregulated.

## Discussion

We investigated inflammatory, injury-associated, and molecular responses to two concentrations of ozone in adult, female C57BL/6J mice. The design of this study facilitated examination of gene expression responses specific to and shared across two key target tissues, the conducting airways (CA) and airway macrophages (AM), along with identifying concentration-specific transcriptional patterns.

We first used a principal components analysis (PCA) to explore our gene expression data within each compartment and gain large-scale insights into the structure of our data. Our PCA plots demonstrate that separation in gene expression between samples can largely be explained by exposure group, and in the case of CA, mirror concentration-response patterns seen in other phenotypes such as airway neutrophilia. We then conducted differential gene expression analysis, identifying which genes were similarly up- or down regulated in each compartment as a function of O_3_ concentration. Overall, CA tissue had a more dynamic transcriptional response to O_3_ exposure than AM as judged by the number of DEGs when comparing 1 ppm or 2 ppm versus FA exposure, though AM had more DEGs when comparing 2 versus 1 ppm exposure. These observed differences may be attributable to the heterogeneous cellular composition of conducting airway tissue compared to airway macrophages.

Previous studies of airway epithelial responses to O_3_ exposure have identified the importance of epithelial-derived immune signaling and oxidative stress responses (DEVLIN *et al*. 1994; WANG *et al*. 2006). Our results confirm that genes critical for these processes (e.g., *Krt6a*, *Orc1*, *Gsta1*, *Mt2*) are upregulated in CA shortly after O_3_ exposure, along with enrichment of pathways associated with tissue regeneration and cellular proliferation. Together, this supports the findings of our histological analyses: namely, the airways were denuded, which increased in severity in a concentration-dependent manner. Though many immune-related genes were upregulated after O_3_ exposure, we also identified a number of genes that were down regulated including *Cxcr3*, *Ifitm1*, *Ccr9*, and *Il12rb*.

AM are known to have dual pathogenic and protective roles after O_3_ exposure, including secretion of both pro- and anti-inflammatory mediators (PENDINO *et al*. 1995; SUNIL 2012) and scavenging of cellular debris (ARREDOUANI *et al*. 2007). O_3_ exposure is also associated with AM dysfunction, including decreased phagocytic and efferocytic function (GILMOUR *et al*. 1991; JAKAB *et al*. 1995), and increased susceptibility to respiratory infections (MIKEROV 2008). Thus, identifying molecular mechanisms that drive these functional outcomes is critical. Many of the most significantly upregulated DEGs were involved in immune signaling including genes encoding chemo- and cytokines (e.g., *Ccl24*, *Cxcl3*, *Ccl17*) and canonical markers of macrophage polarization (e.g., *Retnla*, *Arg1*, *Cd80*), consistent with the findings of previous studies (LASKIN *et al*. 2011; SUNIL 2012; LASKIN *et al*. 2019). Significantly down regulated genes included MHC II genes (*H2-M2*, *H2-DMa*, *H2-Q10*), various membrane transporters (*Abcb1a*, *Slc2a5*), and immune modulators (*Ifnb1*, *Nfkbid*, *Tcim*).

By and large, CA and AM exhibited unique transcriptional responses to O_3_. However, we also identified genes that were concordantly upregulated or down regulated in both CA and AM samples. Genes that were upregulated in both compartments included those involved in DNA replication and cell cycle progression like cyclin family members and kinetochore components (*Cdc45*, *Cdk18*, *Cenpk*), proteases and anti-proteases (*Timp1*, *Ctse*, *Prss27*) which might regulate tissue regeneration and repair, and *Spp1* (osteopontin), a gene that was previously identified through a transcriptomic analysis of human O_3_ responses (LEROY *et al*. 2015). A smaller proportion of genes were down regulated in both compartments and spanned a wide range of categories including kinases (*Nek3*), actin- and microtubule-associated proteins (*Espn*, *Tppp3*, *Dnah2*), and chromatin-associated enzymes (*Usp11*, *Hr*).

Using a set of filtering criteria, we were able to categorize the lists of DEGs in each of the two tissue compartments into four broad categories: genes that follow a linear concentration-response, genes that exhibited threshold-like effects, and genes that are only differentially expressed after 1 ppm O_3_ exposure (peak/trough). Overall, a similar number of DEGs from CA and AM could be categorized, though the relative proportions of each of the categories differed. A greater number of CA DEGs were categorized as negative monotonic and negative or positive threshold A (threshold at 1 and 2 ppm). In contrast, more AM DEGs were classified as positive monotonic, positive threshold B (threshold across FA and 1 ppm) and trough. We also examined pathway enrichment and functional annotations associated with specific categories of genes. Within CA, the top pathway-expression pattern pairs were protease activity (positive monotonic), many cell division-associated pathways (peak), and cellular defense responses (negative threshold B). Similarly, a few pathway-expression category pairs were noteworthy in AM including cellular response to heat (trough), DNA damage response (positive threshold A), and cytokine-cytokine receptor interaction (positive monotonic).

Using previously published data from three transcriptomic studies of whole lung responses to O_3_ exposure, we conducted a simple meta-analysis to identify genes that are consistently altered in response to O_3_ exposure. Though no genes were represented across all three previously published studies and our own, five DEGs in CA (*Cdk1*, *Lcn2*, *Mt1*, *Saa3*, *Serpina3n*) and four DEGs in AM (*Cdk1*, *Lcn2*, *S100a9*, *Saa3*) were represented in two studies and our own. We also used these data to demonstrate that whole lung gene expression analyses tend to reflect CA-derived gene expression more clearly than AM-derived gene expression, though a small number of genes differentially expressed only in AM samples were also captured in the whole lung data (e.g., *Ccl17*, *Hmox1*, *Mmp12*, *S100a8*).

Within CA, many genes were differentially expressed that have not been previously associated with O_3_ responses, including *Oxtr*, *Fgf23*, and *Ptx3*. *Oxtr* (oxytocin receptor) was significantly down regulated after both 1 and 2 ppm O_3_ exposure versus FA (1 ppm and 2 ppm FC: 0.125). Down regulation of *Oxtr* has been associated with behavioral changes after diesel exhaust exposure in mice (WIN-SHWE 2014), and may bind to RAGE (receptor for advanced glycation endproducts) (YAMAMOTO 2019), a receptor involved in recognizing damage-associated molecular patterns (DAMPs) that has been implicated in inflammatory responses to acute lung injury (GRIFFITHS AND MCAULEY 2008; BLONDONNET *et al*. 2017). *Fgf23* (fibroblast growth factor 23) followed a concentration-response expression pattern after both 1 and 2 ppm O_3_ exposure (1 ppm FC: 14.9, 2 ppm FC: 48.5), and has been shown to promote airway inflammation and release of IL-1β, in the context of chronic obstructive pulmonary disease (KRICK *et al*. 2018). Finally, *Ptx3,* the gene encoding pentraxin 3, was highly upregulated after 2 ppm O_3_ exposure (FC 17.2). This gene belongs to a family of widely evolutionarily conserved proteins called pentraxins, among which are the acute phase proteins C-reactive protein (CRP) and its murine cognate serum amyloid component P (SAP) (DONI 2019). *Ptx3* is less well characterized, but is induced by pro-inflammatory cytokines such as TNF, along with regulating chronic inflammation and responses to sterile tissue damage (KUNES *et al*. 2012). Though *Ptx3* is known to be expressed by innate immune cells, including macrophages (IMAMURA *et al*. 2007), we note that *Ptx3* was selectively expressed by CA in our study suggesting that it may have specific roles in epithelial responses to O_3_ exposure.

Likewise, we identified a number of novel DEGs in AM after O_3_ exposure including *Gucy2c*, *Hamp*, and *Hes1*. *Gucy2c* (guanylate cyclase 2C) is an enzyme that is typically found in the intestinal epithelium, and is known to bind the endogenous hormones guanylin and uroguanylin to drive intestinal cell differentiation, in addition to binding bacteria-derived diarrheagenic, heat-stable toxins. *Gucy2c* is not known to be expressed in the lung; however, it was highly differentially expressed in AM after 1 ppm O_3_ exposure, and the most highly differentially expressed gene in AM after 2 ppm O_3_ exposure (1 ppm FC: 147, 2 ppm FC: 477.7). *Hamp* (hepcidin) is a hormone primarily produced in the liver (though also produced by lung macrophages(NGUYEN *et al*. 2006)) and operates in a positive feedback loop with circulating iron to regulate serum iron homeostasis. Interestingly, *Hamp* is known to be upregulated in the context of infection and inflammation, but was highly down regulated after 2 ppm O_3_ exposure in our study (FC 0.067). *Hes1* (hairy and enhancer of split 1) was significantly down regulated in AM after 1 ppm O_3_ exposure (FC 0.134), but not after 2 ppm O_3_ exposure. *Hes1* is a repressor gene known to negatively regulate neutrophilic inflammation through suppression of transcriptional elongation of various macrophage-derived chemokine genes, including CXCL1 (SHANG 2016).

While the methods we used enabled us to generally describe the responses of two important lung compartments to O_3_, our approach is not without its limitations. Both tissue compartments we analyzed are heterogeneous, which cannot be easily accounted for using bulk RNA-seq. Flow sorting or single-cell RNA-seq could be used to assign AM origin and phenotype, particularly after O_3_ exposure when inflammatory monocytes have been recruited from the periphery (FRANCIS *et al*. 2017a; FRANCIS *et al*. 2017b). Likewise, these approaches could be applied for the many cell types that comprise the airways and resident and/or recruited immune cell populations that are tightly associated with or interdigitated within the epithelium. This approach would also address the challenge of interpreting whether gene expression in isolated airways of O_3_-exposed mice is primarily due to altered transcriptional activity or a result of changes in cell composition. We also opted to use only female mice, and sex is known to have an impact on gene expression responses in many contexts, including after O_3_ exposure (CABELLO *et al*. 2015; MISHRA *et al*. 2016; FUENTES *et al*. 2018). Thus, additional studies will be required to determine the impact of sex on transcriptional responses in the airway epithelium and macrophages. Nevertheless, our study provides what we believe is the first transcriptomic analysis of responses to O_3_ in murine conducting airways and airway macrophages after whole body exposure.

In summary, we have identified genes that change in the airway after O_3_ exposure in parallel to concentration-dependent increases in airway neutrophilia, pro-inflammatory cytokine secretion, and epithelial tissue injury. Additionally, we demonstrated that the airway epithelium and airway macrophages have both common and distinct transcriptional signatures following O_3_ exposure. The altered genes and pathways presented in our study increase our understanding of the molecular mechanisms that underlie respiratory toxicity to O_3_, and provide candidate genes and pathways for focused future study.

## Supporting information

Description of Supplement

## Acknowledgements

The authors would like to acknowledge the assistance of Kathryn McFadden (laboratory animal support), Michala Patterson (conducting airway microdissection and RNA isolation), Carlton Anderson (UNC Advanced Analytics Core, Luminex processing), the UNC High-Throughput Sequencing Facility (library preparation and RNA-seq), and Courtney Nesline and the UNC Division of Comparative Medicine.

