## Supplementary material for "Transcriptional profiling of the murine airway response to acute ozone exposure": Description of Supplement

**Supplement and Data Details**

Supplemental Table 1. STAR and Salmon mapping statistics for all samples.

Supplemental Table 2. Trend assignments for differentially expressed genes.

Supplemental Table 3. Enrichr analysis of a subset of categorized genes from Supplemental Table 2.

Supplemental Table 4. Overlapping genes between conducting airways and airway macrophages between each treatment comparison.

Supplemental Table 5. Details of the previous three studies used for hypergeomeric enrichment analysis.

Supplemental Table 6. Sets of differentially expressed genes from previous transcriptomic studies of whole lung responses to ozone exposure that were used for hypergeometric enrichment analysis and vote-counting meta-analysis.

Supplemental Table 7. Hypergeometric enrichment analysis of previous transcriptomic studies of lung responses to ozone compared to our genes.

Supplemental Table 8. Vote-counting meta-analysis of genes across three studies and the current study.

Supplemental Table 9. Genes identified through meta-analysis that were unique to each tissue compartment.

Supplemental Table 10. Enrichr libraries used for Gene Set Variation Analysis (GSVA).

Supplemental Data 1. All differentially expressed genes in conducting airways.

Supplemental Data 2. All differentially expressed genes in airway macrophages.

Supplemental Data 3. All differentially enriched GSVA gene sets in conducting airways.

Supplemental Data 4. All differentially enriched GSVA gene sets in airways macrophages.
